# DENVIS: scalable and high-throughput virtual screening using graph neural networks with atomic and surface protein pocket features

**DOI:** 10.1101/2022.03.17.484710

**Authors:** Agamemnon Krasoulis, Nick Antonopoulos, Vassilis Pitsikalis, Stavros Theodorakis

## Abstract

Computational methods for virtual screening can dramatically accelerate early-stage drug discovery by identifying potential hits for a specified target. Docking algorithms traditionally use physics-based simulations to address this challenge by estimating the binding orientation of a query protein-ligand pair and a corresponding binding affinity score. Over the recent years, classical and modern machine learning architectures have shown potential for outperforming traditional docking algorithms. Nevertheless, most learning-based algorithms still rely on the availability of the protein-ligand complex binding pose, typically estimated via docking simulations, which leads to a severe slowdown of the overall virtual screening process. A family of algorithms processing target information at the amino acid sequence level avoid this requirement, however at the cost of processing protein data at a higher representation level. We introduce deep neural virtual screening (DENVIS), an end-to-end pipeline for virtual screening using graph neural networks (GNNs). By performing experiments on two benchmark databases, we show that our method performs competitively to several docking-based, machine learning-based, and hybrid docking/machine learning-based algorithms. By avoiding the intermediate docking step, DENVIS exhibits several orders of magnitude faster screening times (i.e., higher throughput) than both docking-based and hybrid models. When compared to an amino acid sequence-based machine learning model with comparable screening times, DENVIS achieves dramatically better performance. Some key elements of our approach include protein pocket modelling using a combination of atomic and surface features, the use of model ensembles, and data augmentation via artificial negative sampling during model training. In summary, DENVIS achieves competitive to state-of-the-art virtual screening performance, while offering the potential to scale to billions of molecules using minimal computational resources.

**Graphical TOC Entry:** 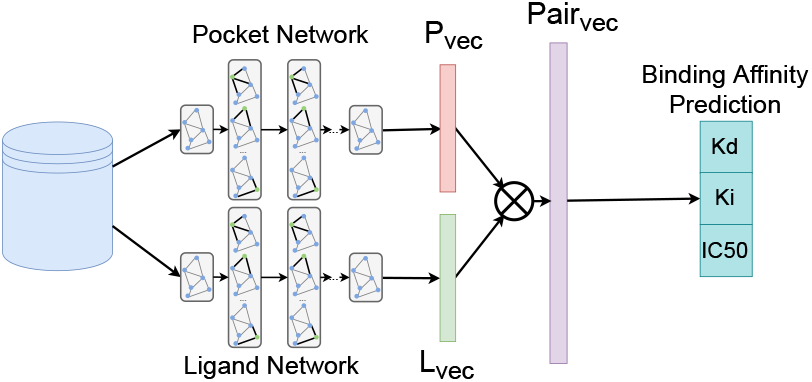

## Introduction

In the pharmaceutical industry, the average cost associated with the development of a new drug is approximately $2.6 billion. At the same time, success rates at the final clinical trial stage are typically below 10%^1^. One of the early phases of drug discovery concerns high-throughput screening, a process by which a large number of chemical or biological substances are tested against a specified target of interest. Machine learning has the potential to substantially accelerate the drug discovery life cycle by performing virtual screening of large databases of candidate molecules against a pharmacological agent (e.g., a protein) associated with some known disease^2^. Virtual screening refers to the process by which existing compounds, potentially with some known properties, are utilised to treat novel diseases^3,4^. The special case where the studied compounds have undergone clinical testing and acquired approval from regulatory bodies (e.g., FDA) is referred to as drug repurposing. The outbreak of the COVID-19 pandemic has revealed an utmost need for developing quick and efficient response strategies against emerging diseases using the virtual screening paradigm. Since the outbreak, an unprecedented global effort has been put into using computational methods to screen vast compound libraries in order to identify candidate molecules for COVID-19 treatment^5^.

Virtual screening can be divided into structure-based and ligand-based. Structure-based virtual screening (SBVS) seeks to capture protein-ligand interactions by using, for example, x-ray crystallography measurements to build models that can generalise to novel target-compound pairs^4,6^. On the other hand, ligand-based virtual screening (LBVS) exploits statistical relationships between ligands that are known to bind to a specific target with the aim of predicting binding properties of novel ligands for the same target^7^.

Traditional approaches for SBVS employ empirical scoring functions and molecular docking simulations to predict binding affinity scores for a target-ligand pair under various docking conformations^8^. Several docking algorithms are currently commercially available^9–12^. Over the last decade, the use of machine learning has been proposed as a means of building upon this approach to achieve higher predictive performance. To that end, a wide variety of machine learning algorithms have been used, ranging from random forests^13^, feed-forward multi-layer perceptrons (MLPs)^14^, convolutional neural networks (CNNs)^15–21^, and graph neural networks (GNNs)^22–25^. Methods based on deep learning are becoming increasingly popular in this domain. For instance, GNINA^16,20,21^ and KDEEP^17^ use three-dimensional (3D) CNNs for binding affinity prediction. The input to both models is a 3D voxel representation of a protein-ligand interaction structure, which is typically estimated by a docking algorithm. In a similar fashion, OnionNet extracts atomic features from 3D representations of protein-ligand complexes, which are then processed by a series of convolutional and fully connected layers to predict binding affinity scores^19^. In addition, methods based on GNNs have been increasingly used in virtual screening applications. The atomic CNN makes use of the 3D protein-ligand complex structure and a specially designed graph convolutional network to predict the binding affinity of the complex^15^. Similarly, Torng and Altman ^23^ employ an amino acid GNN to represent protein pockets and use it in a binding classification task. Finally, Morrone et al. ^25^ use a dual-graph approach to combine protein-ligand docking pose and ligand structure with dedicated GNNs, which are then used in a binding binary classification task.

A limitation of this family of methods is that they typically require either access to the experimental crystal structure^15,17,22,24^ or rely on docking simulations to estimate the protein-ligand binding pose^16,18–20,25^. Docking simulations are typically very slow and therefore greatly decrease the throughput of the overall screening pipeline. For instance, McNutt et al. ^21^ report an average computational time of approximately 25 - 29 s for predicting the binding affinity score of a single protein-ligand pair with GNINA. Unfortunately, the high computational cost associated with docking-based methods can hinder the scalability of the virtual screening pipeline, unless huge computational resources (i.e., several thousands of GPUs) are available^26^.

A separate line of research has investigated the prediction of protein-ligand binding properties by processing target amino acid sequences and ligand two-dimensional structures, for example, using the SMILES representation. This approach ignores the protein and ligand 3D structure altogether. For example, DeepDTA represents protein sequences and ligand SMILES using sets of words that are then processed by CNNs^27^. WideDTA extends this model by including additional textual features in the protein and ligand representations^28^. PADME combines compound and protein sequence composition features into a combined vector and uses a feed-forward MLP to predict binding affinity scores. A variant of the model uses two-dimensional structural information for ligands, but not for proteins^29^. In addition, DeepConv-DTI uses a convolutional layer to process protein sequences and a fully connected layer for drug fingerprint descriptors that are then combined using a fully connected layer to estimate binding affinity scores^30^. DeepAffinity achieves the same objective by using a structural property sequence representation and a combination of convolutional and recurrent layers.^31^ More recently, GNNs algorithms operating on protein sequence data have also been used, including models such as GraphDTA^32^, DGraphDTA^33^ and MONN^34^. While sequence-level models avoid the requirement for docking simulations, this comes at the cost of processing protein information at a higher representation level (amino acid sequence as opposed to atom domain). Despite that promising results have been reported using this family of models, a performance comparison between atom-level and sequence-level algorithms is currently missing from the literature.

In this work, we introduce deep neural virtual screening (DENVIS), a purely machine learning-based, high-throughput, end-to-end-strategy for SBVS using GNNs for binding affinity prediction. Our method does not require access to the experimental protein-ligand crystal structure, neither uses docking simulations to estimate it; hence, it is readily-applicable to real-life situations and also highly-scalable. The main contribution of our work can be summarised as follows:

- development of an end-to-end virtual screening system with highly-competitive performance and extremely fast screening times;
- fusion of multiple protein pocket representations (atomic and 3D surface) in combination with ensemble modelling;
- a data augmentation scheme employing artificial negative sampling during binding affinity network training;
- a systematic benchmark of a wide range of docking-based, machine learning-based, and hybrid docking/machine learning-based algorithms for virtual screening, by employing a cross-database validation strategy^35^ and using an appropriate baseline model^36,37^.

We show that DENVIS achieves competitive, and in some cases better performance than several established methods, including both research and commercial docking algorithms, but exhibits three to four orders of magnitude faster screening times. We additionally show that our method largely outperforms a purely machine learning-based approach with comparable screening times. We showcase the key components of our method by performing extensive ablation studies, and finally discuss implications and directions for future work.

## Methods

### DENVIS high-level overview

We tackle the virtual screening problem via ranking of all possible ligands for each target protein. The ranking is performed using estimates of binding affinity scores for all protein pocket-ligand pairs for a given target. We utilise GNNs to extract high-dimensional, continuous vector representations for ligands and protein pockets separately. These vectors are then combined via an outer product layer and passed onto a regression network predicting multiple binding affinity metrics for each protein pocket-ligand pair (Figure 1(a)).

**Figure 1:**
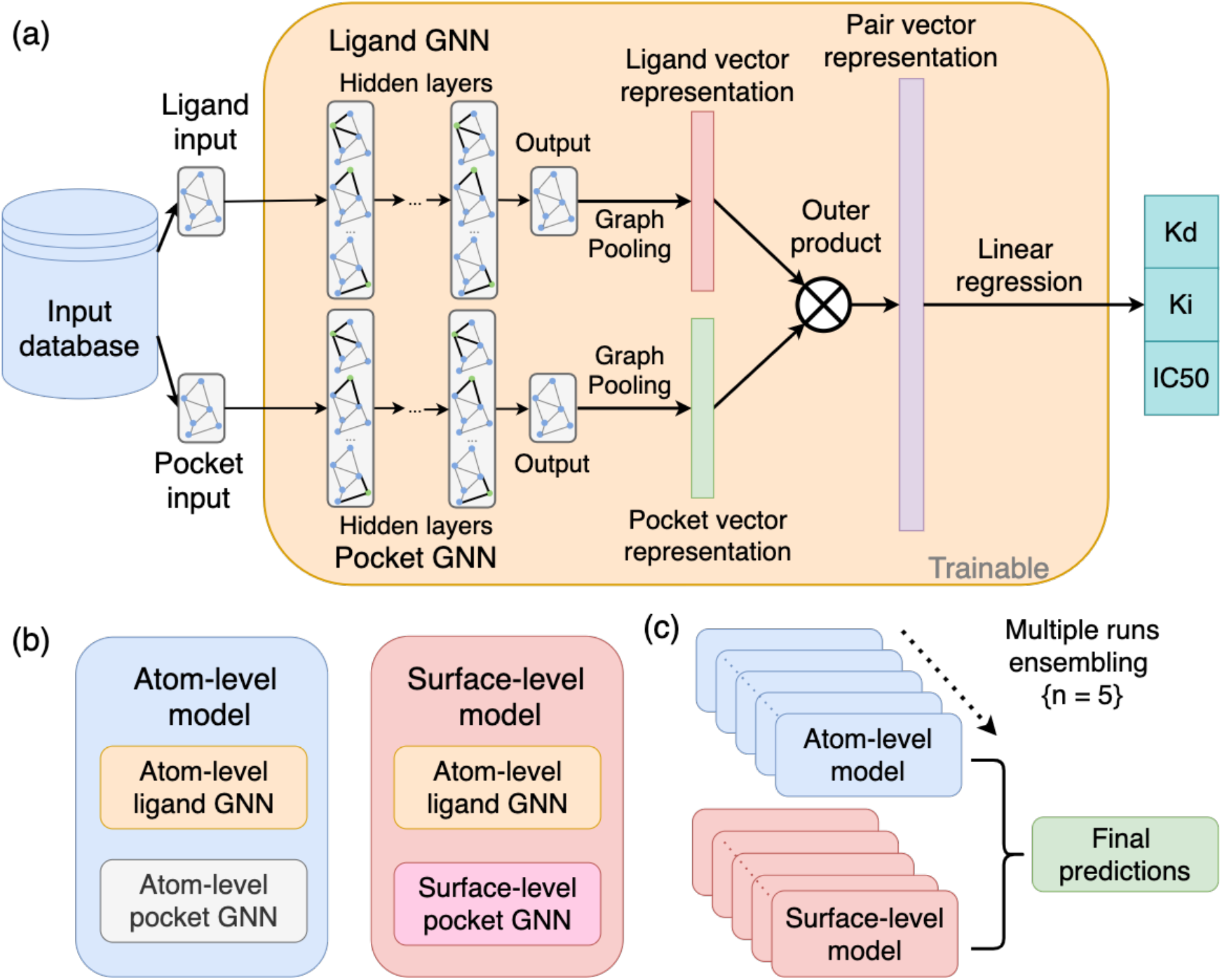
**(a)** DENVIS schematic representation. Ligand and protein pocket features are initially fed to dedicated GNNs. Each GNN yields a graph output which is converted to a vector representation via graph average pooling. The two vectors are then combined via an outer product operation to get a representation for the protein pocket-ligand pair. This final representation is fed to a multi-output linear regression layer to estimate multiple binding affinity metrics. These are *K*_*d*_, *K*_*i*_ and *IC*_50_ when the network is trained on the PDBbind general set, and *K*_*d*_ and *K*_*i*_ when it is trained on the PDBbind refined set. **(b)** Two types of representations and associated GNN models are used for protein pocket data processing: atom-level and surface-level. For ligands, the atom-level approach is used. **(c)** Schematic representation of our combined ensembling strategy for virtrual screening. We train five model instances with different random seeds for both atom- and surface-level approaches. To estimate the final binding affinity prediction score for each type of network, we compute the average scores across the five instances. The final binding affinity prediction scores are computed using a weighted average of the atom- and surface-level ensemble model scores.

We follow two modelling approaches for protein pockets, one based on atomic features and another one based on 3D surface representation. We term these two models *atom-level* and *surface-level*, respectively. The atom-level model consists of a modified version of the graph isomorphism network (GIN)^38^, a generic, yet powerful GNN implementation that has been used in biological and chemical applications^39^. We adopt the feature extraction pipeline of the open graph benchmark^40^ for molecules. The surface-level approach utilises a mixture model network (MoNet)^41^, a specialised GNN with a convolution operation that respects the geometry of the input manifold. We extract chemical and geometrical features from the protein pocket surface mesh^42^. For ligand feature extraction, we follow the atom-level approach (Figure 1(b)).

Our complete pipeline combines predictions of the two protein pocket representation approaches with late fusion. We first train models with the two approaches independently. The final score for a protein pocket-ligand pair is then computed as a weighted average score of the two different models. We additionally use model ensembles, whereby several identical networks are trained independently with different random seeds for each of the atom- and surface-level models (Figure 1(c)).

### Datasets

We utilise several different datasets in various steps of our overall pipeline. Namely, we use TOUGH-M1^44^ for protein pocket and ligand GNN pre-training, the PDBbind v2019 refined and general sets^45^ for binding affinity network training (i.e., training sets), the PDBbind v2019 core set^48^ for model selection (i.e., validation set), and finally the Directory of Useful Decoys: Enhanced (DUD-E) and LIT-PCBA^46^ for model evaluation and benchmarking (i.e., test sets). The characteristics of the various datasets used in this work are summarised in Tables 1 and 2.

**Table 1:** Pre-training and training datasets

| Dataset | # Complexes | # $K_d$ | # $K_i$ | # $IC_{50}$ | Used for | Ref. |
| --- | --- | --- | --- | --- | --- | --- |
| TOUGH-M1 | 7524 | n/a | n/a | n/a | pre-training | <sup>44</sup> |
| PDBbind general set | 17679 | 6455 | 4669 | 6555 | training | <sup>45</sup> |
| PDBbind refined set | 4852 | 2464 | 2388 | 0 | training | <sup>45</sup> |

**Table 2:** Validation and test datasets

| Dataset | # Targets | # Actives/target | # Decoys/target | Used for | Ref. |
| --- | --- | --- | --- | --- | --- |
| PDBbind core set | 57 | 5 | 56 (per active)* | validation | <sup>45</sup> |
| DUD-E | 102 | 40 - 592 (155)† | 50 (per active) | testing | <sup>46</sup> |
| Lit-PCBA | 15 | 13 - 7168 (101))† | 4168 - 362049 (245522))† | testing | <sup>47</sup> |

\* Decoys shared across targets.
† Numbers in brackets indicate medians.

### Pre-training dataset: TOUGH-M1

TOUGH-M1 is a database designed for binding site matching ^44^. It consists of 7,524 distinct pocket structures for which binding ligands are known. These are grouped according to the similarity of their binding ligand resulting in 505,116 pocket pairs with similar binding ligands (positive) and 556,810 pocket pairs with dissimilar ligands (negative). We use this dataset for pre-training protein pocket and ligand GNNs.

### Training and validation datasets: PDBbind

PDBbind is a database with selected entries in the protein data bank (PDB) with protein-ligand complex structures. It comprises complexes for which binding affinity has been experimentally verified using one of three measures: dissociation constant (*K*_*d*_), inhibitor constant (*K*_*i*_), and half maximal inhibitory concentration (*IC*_50_)^49^. The database is updated on a regular basis to include recent additions to PDB and is split into three subsets, namely, general, refined, and core sets, with increasingly stricter criteria in data quality for inclusion. Among them, only the general set includes complexes with *IC*_50_ affinity measurements. In this work, we use v. 2019 of the database^45^, which comprises 17,679, 4,852, and 285 complexes in the general, refined, and core sets, respectively. We experiment with both the refined and general sets for model training and use the core set as a validation set for model selection. To avoid overlap in the raw data of training and validation sets, we filter out the core set entries from both the general and refined sets. When using the general set, we only consider protein-ligand complexes and completely discard other types of structures (e.g., protein-protein complexes).

### Test datasets: DUD-E and LIT-PCBA

We use the DUD-E database^46^ as the main virtual screening benchmark in our study, since it is widely used and there exist available prediction outputs for a variety of models^9–14,21,50^. We additionally validate our method on the recently published LIT-PCBA benchmark^47^, for which we have access to prediction outputs from one competitive method^50^.

The DUD-E^46^ consists of 102 protein targets with an average of 224 active ligands per target. For each active ligand, there are 50 decoys (i.e., inactive compounds) selected from the ZINC database^51^ with similar physical properties but different chemical structures, which are considered to be non-binders. This results in over a million protein-ligand pairs that can be used for screening evaluations. No binding affinity measurements are provided with this dataset.

LIT-PCBA^47^ is a recently released virtual screening dataset that contains 15 protein targets with 9780 distinct actives and 407839 different inactives selected from PubChem. Many of the inactives are shared between targets which results in more than 2.5 million protein ligand pairs. Additionally, multiple experimental structures / templates are included for 13 out of 15 targets. The database mimics experimental screening decks with respect to hit rates and potency distributions. Special care has been taken to include topologically similar active and inactive compounds. Furthermore, all compounds for each target–both actives and inactives–are taken from single assay data, and therefore there is experimental support for both activity and inactivity. As with DUD-E, no binding affinity measurements are provided for the active compounds. The authors of the database propose fixed training and validation splits. We do not make use of these splits, but instead we evaluate the performance of the models on the full dataset.

### Preprocessing and feature extraction

The various datasets that we use have been processed using slightly different pipelines. To avoid systematic discrepancies arising from differences in preprocessing steps and ensure homogeneity across datasets, we perform a common preprocessing pipeline for all protein structures. Firstly, we download the raw protein data from a common source, namely the PDB. For the specific datasets that we use, protein structures in the PDB are typically available as part of a protein-ligand complex. We then protonate the protein data^49^ and remove any atom that is not part of the protein. We perform all pre-processing steps using the rdkit library (v2021.09.2)

Similarly, we follow a consistent methodology for pocket extraction to avoid differences across datasets. It should be noted that our modelling approach is agnostic to the used pocket extraction method. Since in our case we have access to protein-ligand complexes, we follow the the PDBbind database creation procedure and keep the atoms of the protein within a 10 Å radius from the bound ligand^45^. In cases where no complex structure is available, the pocket can be extracted using any binding site detection algorithm, such as Fpocket^52^.

As explained earlier, and also illustrated in Figure 2, we process protein pocket information using two distinct levels of representation: atom-level and protein surface-level. For ligand feature extraction, we employ the atom-level approach. The following sections describe the two feature extraction pipelines.

**Figure 2:**
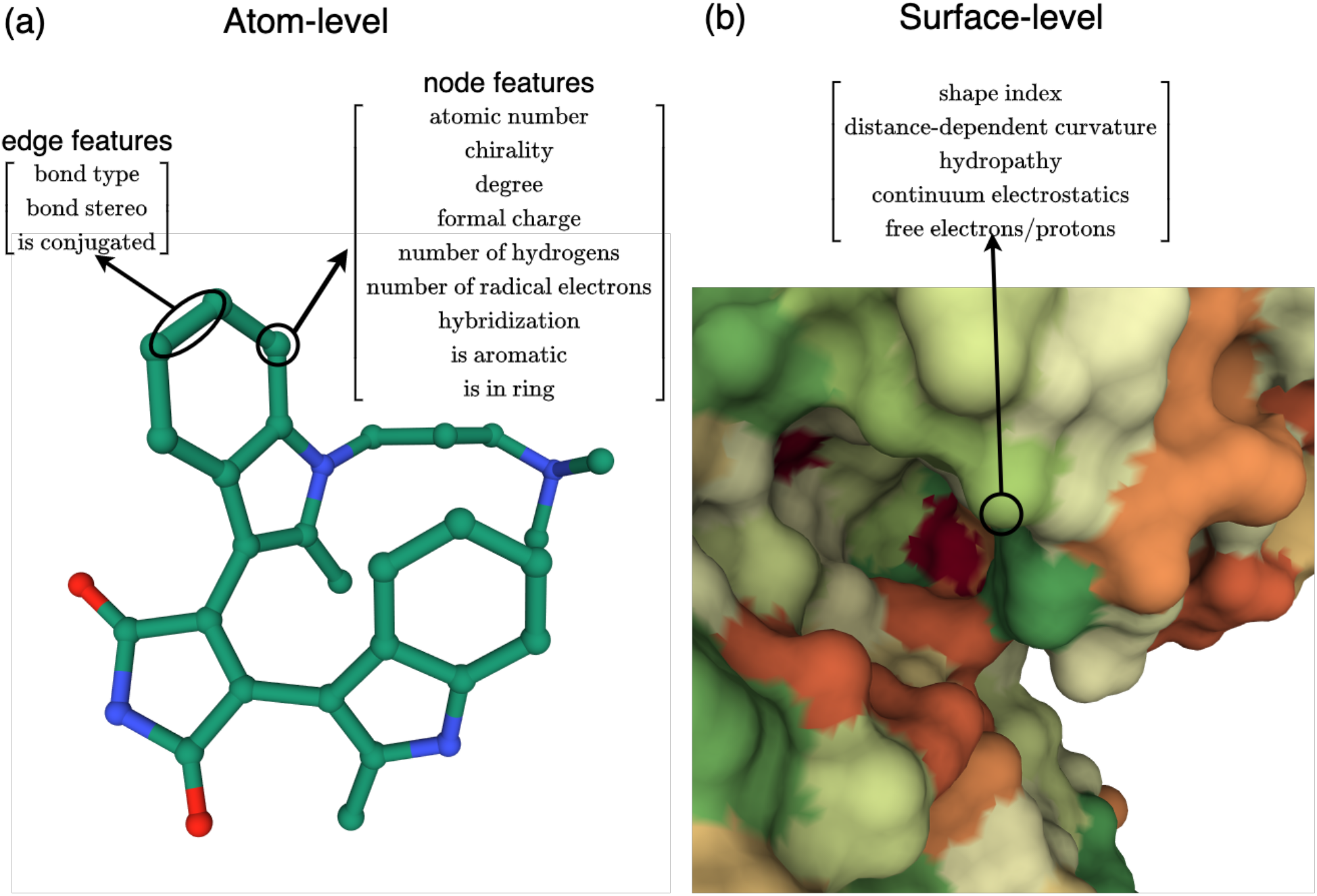
Feature extraction from multiple representations. (a) Atom-level approach. Nine node and three edge categorical features are extracted from the raw structural data, for both protein pockets and ligands^40^. (b) Surface-level approach. Two geometrical and three chemical features are extracted from the protein pocket surface^42^. Visualisations are created using the Mol* tool^43^, available on the PDB website, and correspond to the human protein kinase C beta II complexed with a bisindolylmaleimide inhibitor (PDB code: 2i0e).

### Atom-level feature extraction

In the atom domain, we extract nine node and three edge features for each molecule (protein pocket or ligand)^40^. The node features include the atomic number, chirality tag, degree, formal charge, number of explicit hydrogens, number of radical electrons, hybridisation, and two binary features indicating whether the atom is aromatic and whether it is in the ring. The edge features include the bond type, bond stereochemistry, and a binary feature indicating whether the bond is conjugated (Figure 2(a)). All atomic node and edge features as treated as categorical variables.

### Surface-level feature extraction

With our surface-level approach, we extract both geometrical and chemical features for the protein pocket surface^42^. Geometrical features consist of the shape index and the distance-dependent curvature, which is an 8D vector. Chemical features include hydropathy, continuuum electrostatics, and free electrons/protons (Figure 2(b)). We then keep the binding interface defined as the surface nodes that are within a 3 Å radius from the bound ligand. Note that in this case, graph nodes correspond to vertices on the extracted 3D surface mesh, as opposed to the atom-level case, where the nodes correspond to atoms in the molecular structure.

In some complexes, ligands are bound in the interior of the protein and, as such, do not interact with the protein surface. Due to a limitation of the surface extraction routine^42^ preprocessing is impossible in this case, and thus we omit those proteins from our surface-level models. This results in eight out of 102 targets from the DUD-E dataset being discarded. For these targets we base our predictions entirely on the atom-level models. For surface-level model training, we discard 691/17679 and 256/4852 complexes from the PDBbind general and refined sets, respectively, due to issues related with pocket surface extraction.

### Graph neural network architectures

We use two types of GNNs processing information at the two levels of representation: atom-level and protein surface-level. The atom-level GNN is used for both protein and ligand inputs, whereas the surface-level GNN is only used for proteins.

#### Atom-level model

For the atom-level GNN, we use our own modified version of the GIN layer^38^, which additionally supports edge feature updates at each layer. Briefly, at each GIN layer the features of every node are summed with the corresponding features of the neighbouring nodes and fed into a shallow feed-forward network (i.e., two-layer MLP with ReLU activation), which computes the updated node features. In parallel, the features of each node and the summed features of its neighbours are concatenated with the edge features and fed into a separate, yet identical, MLP to compute the updated edge features.

All atom and edge features are categorical variables^40^. Thus, the first component of the atom-level GNN comprises a collection of embedding layers that convert the categorical variables into continuous vectors. The output size of each embedding layer is computed using the following empirical rule^53^:

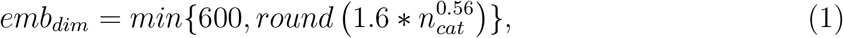

where *emb*_*dim*_ denotes the embedding layer output dimensionality, and *n*_*cat*_ is the cardinality of the categorical variable. The outputs of all embedding layers are concatenated along the feature dimension. Using the empirical rule in Equation 1, the dimension of the continuous node and edge feature vectors are *d*_*node*_ = 56 and *d*_*edge*_ = 10, respectively. The node and edge feature vectors are then fed to a series of five blocks comprising a GIN layer, a batch normalisation layer, a ReLU activation function, and a dropout layer. After the final block, where the activation function is omitted, a graph vector representation is computed using a graph average pooling operation. The node feature dimensionality is progressively increased at each layer in a linear fashion, such that the final vectors have dimensionality *d*_*emb*_ = 300.

#### Surface-level model

For our surface level model, we use the MoNet architecture, which is a specialised network that performs a geodesic convolution on the protein surface with learnable gaussian kernels^41^. Briefly, the filters on a geodesic convolution are calculated with respect to distances between points on the surface manifold, as opposed to distances in a Euclidean space. The surface input consists of chemical and geometric features extracted from the patch that is adjacent to the bound ligand. We use a 2 layer MoNet with output dimensionality *d*_*emb*_ = 32^42^. As with the atom-level GNN, the final protein vector representation is obtained with a graph average pooling operation.

### Graph neural network pre-training

We use the TOUGH-M1 dataset to pre-train three separate GNNs: 1) atom-level protein pocket GNN; 2) surface-level protein pocket GNN; and 3) atom-level ligand GNN. The three pre-trained networks are then used to initialise the weights of the respective components of the binding affinity networks described below. The pre-training procedure is performed independently for each of the three GNNs. Our motivation for employing network pre-training stems from the fact that the number of labelled samples in the training set (i.e., PDBbind) is rather limited (in the order of a few thousand). Therefore, our goal is to obtain a good model parameter initialisation for binding affinity network training (i.e., fine-tuning).

Since the TOUGH-M1 dataset is organised in similar/dissimilar pairs of protein pockets and, additionally, similar/dissimilar pairs of ligands, we follow a supervised metric learning approach based on Siamese networks^54^. It should be emphasised that this training procedure is fundamentally different to the one used for binding affinity network training, in which case input data come in protein pocket-ligand pairs and the labels are the corresponding binding affinity measurements. In contrast, during network pre-training, input data come in protein pocket pairs, in the case of protein pocket GNNs, or ligand pairs, in the case of the ligand GNN. The target labels are in this case binary, indicating whether two molecules (protein pockets or ligands, depending on which GNN is being pre-trained) are similar or dissimilar. During pre-training, the two feature vectors corresponding to the two molecules in the pair are fed to the same GNN to compute the corresponding embedding vectors, which are then used to compute the Euclidean distance between the molecules in the embedding space. The objective of the training procedure is to enforce pairs of similar molecules to be closer in embedding space (i.e., have smaller distance) than pairs of dissimilar ones. To that end, we use the normalised temperature-scaled cross-entropy (NTXEnt) loss function^55^.

We adopt a transitive similarity approach to identify similar/dissimilar pairs within a batch. That is, we label as similar/dissimilar all possible pairs that can be inferred from the batch, as opposed to using only the ones that are explicitly defined. Consider, for example, a batch with two pairs (A, B) and (B, C), with corresponding labels “similar” and “dissimilar”. By using the transitive similarity approach, the pair (A, C) will be also labelled as “dissimilar” and will be used in the computation of the loss function. For model pre-training we use an Adam optimiser with learning rate of 0.001, a batch size of 8, and train for 10 epochs. We set the value of the temperature hyper-parameter of NTXEnt loss to 0.07.

### Binding affinity prediction networks

A high-level overview of our system is shown in Figure 1(a). In this section we provide the details of our implementation and the training procedure.

We use the PDBbind dataset to train our binding affinity prediction networks and experiment with both the refined and general sets for training. All binding affinity measurements are converted to negative log-scale using a base of 10, that is, we perform the transformation *pK* = *−* log_10_*{K*_*d*_, *K*_*i*_, *IC*_50_*}*. We use all protein pocket-ligand pairs from the training datasets (general or refined) and refer to them as *positive* examples. Moreover, we generate artificial *negative* examples during training by combining protein pockets and ligands from different complexes and assigning them a negative log-binding affinity score of 0. That is, we make the following assumptions: 1) protein pockets and ligands from different complexes do not bind; and 2) for non-binding protein pocket-ligand pairs the corresponding binding affinity score is *−∞*. A different set of negative examples is sampled at the start of every training epoch, while the ratio of positive/negative examples is fixed to unity throughout training.

As noted in the previous sections, we use two distinct representations for target protein pockets. Hence, we train two networks independently, one using atomic features and a separate one using 3D surface features. We term these two networks *atom-level* and *surface-level* models, respectively. Each model comprises two sub-networks encoding protein pocket and ligand features into continuous vectors, which are initialised using the pre-training procedure described in the previous section. The protein pocket and ligand vector representations are then combined using an outer product operation. The resulting pair vector represntation is fed to a dropout layer and finally to a fully connected linear layer with multiple outputs, each one corresponding to one binding affinity measure. These are *K*_*d*_, *K*_*i*_, and *IC*_50_ when the networks are trained on the PDBbind general set, and *K*_*d*_, *K*_*i*_ when they are trained on the PDBbind refined set (Table 1). Our design choice for a multi-output network is based on: 1) biological intuition regarding the different nature of the three binding affinity metrics, especially between *IC*_50_ and the other two metrics^56^; and 2) the qualitative observation that these follow different distributions (Supporting Figure S1). We observe that the three binding affinity metrics have similar ranges and, thus, assign an equal weight to the loss term associated with each network output. The overall loss is then computed as:

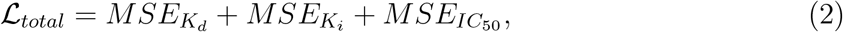

where *MSE* denotes mean squared error, and the *MSE*_*IC*50_ term is omitted in the case of training on the PDBbind refined set.

For our ensembling modelling approach (Figure 1(b)), we train multiple model instances using different random seeds for negative sample augmentation (and weight initialisation in one of our ablation studies). Based on preliminary analysis using the PDBbind core set as a validation set, we find that performance plateaus at around *M* = 5, where *M* denotes the number of model instances in the ensemble (Figure S3 in the Supporting Information (SI)). Thus, we choose to use five models for each of the atom- and surface-level ensembles, as this offers a good trade-off between performance and inference time, which scales linearly with *M*.

Unless noted otherwise, we train our networks for 600 epochs using the Adam optimiser with learning rate of 0.001 and a batch size of 8. We set the weight decay to 0.001 and dropout probability of all layers to 0.3.

### Virtual screening strategy

Once our binding affinity networks have been trained, we use them to perform virtual screening. As described in the previous section, our networks have either two or three outputs corresponding to the binding affinity measures available in the training dataset(s). For a specified network and protein-ligand pair, the final predicted affinity score is computed as the unweighted average of the outputs of the network. For each of the two network types (atom-level and surface-level), we run inference on the query protein-ligand pair separately for each of the five model instances and compute the unweighted average scores across the different models (i.e., *multiple runs ensembles*). Finally, we compute a weighted average score of the atom-level and surface-level models. In mathematical terms, the predicted negative log binding affinity score *ŷ* (unitless) for a protein pocket-ligand pair (*p, l*) is estimated as follows:

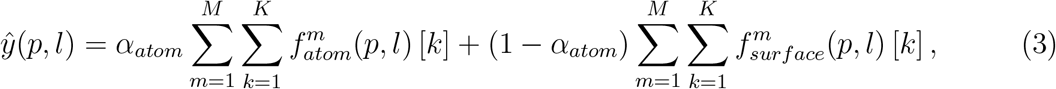

where 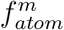 and 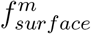 denote the *m*^th^ instance of the atom-level and surface-level models, respectively, *k* indexes the outputs of each of the models, *K* = 3 is the number of model outputs (or *K* = 2 for models trained on the PDBbind refined set), *M* = 5 is the number of model instances trained for each configuration, and *α*_*atom*_ is the weight coefficient for the atom-level model, which is set *a priori* to *α*_*atom*_ = 0.5. For targets for which surface preprocessing is not successful, we set *α*_*atom*_ = 1.0 (eight out of 102 DUD-E targets).

Once binding affinity scores have been estimated using Equation 3, ligands are ranked in decreasing binding affinity prediction order on a per-target basis. The resulting per-target ligand rankings are then used to compute the three performance scores introduced in a later section.

#### Efficient virtual screening implementation

We implement an efficient screening strategy for our method. That is, we first compute the vector representations for each protein pocket and ligand in the screening dataset and store them in memory. Then, for each protein pocket-ligand pair query, we retrieve the corresponding vectors from memory and feed them to the interaction outer product and final linear layers. This enables us to reduce inference times, since most computation is spent on extracting the vector representations from the protein pocket and ligand graph data. This efficient strategy can be adapted, if required, such that only target protein vector representations are pre-computed and stored in memory. This is useful in cases where the screening dataset has a small number of targets, but a very large number of ligands that cannot fit in memory (e.g. DUD-E). Even in this case, pre-computing protein pocket vectors can be very efficient as compared to the naive case where protein pocket vectors are computed afresh, due to the comparatively larger size of protein pocket graphs, which leads to a higher computational cost for the protein pocket GNN.

### Evaluation

#### Performance metrics

We use three metrics for performance evaluation: area under receiver operating characteristic curve (AUROC), enrichment factor (EF) and Boltzmann enhanced discrimination of ROC (BEDROC). All three metrics are computed separately for each target protein and, unless noted otherwise, median scores across targets are reported (i.e., *micro-averaging*). To compute AUROC and BEDROC scores for a given target we use the sorted list of predicted binding affinity scores. It should be noted that although these scores do not correspond to binding probabilities, they can still be used for the computation of the two metrics.

The use of AUROC has been often criticised in virtual screening applications, as this metric assigns equal weight to all ligands in the screening library without taking into account their ranking. Thus, the metric does not take into consideration the fact that only a small fraction of top-ranked ligands will be experimentally validated (i.e., *early recognition*). To address this issue, BEDROC uses exponential weighting to assign larger weights to early rankings^57^ as follows:

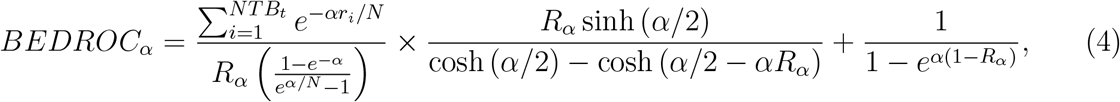

where *NTB*_*t*_ is the total number of true binders (actives), *N* is the total number of ligands, *R*_*α*_ = *NTB*_*t*_*/N* is the ratio of actives in the screening library, and *r*_*i*_ is the rank of the *i*th active moleclue. We set *α* = 80.5, which results in the 2% top-ranked ligands accounting for 80% of the BEDROC score^58^.

The EF is another screening power metric which evaluates the ability of a screening algorithm to identify true binders among the top-ranked ligands^59^. It is related to the widely used recall metric, but only considers a subset of top-ranked ligands. The EF metric is defined as follows:

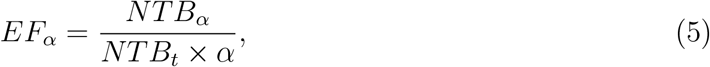

where *NTB*_*α*_ is the number of true binders in the top *α*% (e.g., *α* = 1%, 5%, or 10%) and, as above, *NTB*_*t*_ is the total number of true binders in the screening library. We choose *α* = 1 and thus evaluate recall among the top 1% of our ranking list.

#### Benchmark

We use the DUD-E and LIT-PCBA databases as benchmarks to compare the virtual screening performance of DENVIS against several competitive methods, falling into three main categories: 1) docking algorithms (AutoDock Vina^60^, Gold,^9^ Glide,^11^ FlexX,^12^ Surflex^10^); 2) hybrid docking/machine learning-based algorithms (GNINA^21^, RF-score^13^, NN-score^14^); and 3) a purely machine learning-based algorithm (DeepDTA^27^). Note that GNINA, RF-score, and NN-score require the binding pose of the query protein-ligand pair and for this purpose they make use of AutoDock Vina binding pose predictions. GNINA uses a 3D CNN applied to the docked protein-ligand pose, whereas RF-score and NN-score improve upon AutoDock Vina binding affinity estimates by using a random forest and a shallow feed-forward network, respectively.

For the four commercial docking methods (i.e., Gold, Glide, FlexX, and Surflex), we use the predictions from Chaput et al. ^58^ (DUD-E). For details on the configurations used for each of these algorithms we refer the reader to the original publications^9–12,61–64^. For AutoDock Vina, RF-score, and NN-score, we use the results from Ragoza et al. ^16^ (DUD-E). Finally, for GNINA, we use the results reported by McNutt et al. ^21^ (DUD-E) and Sunseri and Koes ^50^ (LIT-PCBA).

To compute binding affinity predictions with DeepDTA, we make use of the DeepPurpose library^65^. In contrast with all other methods in the benchmark, which make use of structural protein and ligand information, DeepDTA is a sequence-level model and makes predictions by processing the target amino acid sequence and SMILES strings. We experiment both with the model provided by DeepPurpose which, as originally proposed by the authors is pre-trained on the DAVIS dataset^66^, as well as our own version of the model that we train from scratch on the PDBbind refined set. We use a learning rate of 10^*−*4^, a batch size of 256, and train the network for 600 epochs.

#### Ligand baseline

We additionally assess the performance of a simple ligand baseline model. To that end, we train binding affinity networks by removing the protein pocket GNN altogeher. That is, the model learns to assign binding affinity scores to ligands without considering any type of protein information. Such ligand-based baselines have been previously proposed as a means of quantifying potential dataset biases^36,67^. To avoid considering the same inputs as both positive and negative examples, we switch off negative sampling in this case during training.

#### Statistical analysis

The DUD-E database comprises 102 targets, which makes it possible to run statistical comparisons for our benchmark with acceptable statistical power. Without making any assumptions about the underlying distribution of AUROC, EF_1_, and BEDROC_80.5_ scores, we use the Friedman test to assess the effect of decoding algorithm on performance. We perform post-hoc pair-wise comparisons using the Wilcoxon signed-rank test and the Šidák correction method to account for multiple comparisons. For LIT-PCBA, the number of targets is much smaller (i.e., 15), which would translate into prohibitively small statistical power. For this reason, we omit to run statistical analyses on this dataset.

## Results

### Virtual screening performance benchmark: DUD-E

We train five instances of atom- and surface-level models using the PDBbind refined and general sets (v. 2019). Model training dynamics for the two types of models and two training datasets are shown in Figure S2 in the SI. We run inference using all 10 models on the DUD-E dataset, which comprises 102 targets, and combine their predictions to obtain final predictions (Figure 1(c) and Equation 3). We refer to our ensemble models trained on PDBbind general and refined sets as “DENVIS-G” and “DENVIS-R”, respectively. We compare the performance of our ensemble models to all other models in the benchmark and present the overall results in Figure 3. Median scores achieved with all methods are also summarised in Table 3. The Friedman test reveals a significant effect of decoding algorithm on performance (AUROC, *p <* 10^*−*108^; EF_1_, *p <* 10^*−*120^; BEDROC_1_, *p <* 10^*−*120^). We perform post-hoc pair-wise comparisons for these three metrics on a subset of model pairs of interest and present the results in Table 4.

**Table 3:** DUD-E virtual screening benchmark summary (median scores)

|  | AUROC | EF<sub>1</sub> | BEDROC<sub>80.5</sub> |
| --- | --- | --- | --- |
| DENVIS-G<sup>*, †, ∇</sup> | **0.92** | **38.92** | **0.66** |
| DENVIS-R<sup>*, †, ∇</sup> | 0.86 | 18.37 | 0.33 |
| DeepDTA<sup>◦, ∇</sup> | 0.58 | 1.62 | 0.04 |
| Gold<sup>*, ◊</sup> | 0.86 | 21.08 | 0.39 |
| Glide<sup>*, ◊</sup> | 0.82 | 21.96 | 0.35 |
| Surflex<sup>*, ◊</sup> | 0.75 | 9.36 | 0.19 |
| Flexx<sup>*, ◊</sup> | 0.76 | 10.10 | 0.20 |
| Vina<sup>*, ◊</sup> | 0.75 | 6.84 | 0.14 |
| GNINA<sup>*, ◊, ∇</sup> | 0.79 | 15.48 | 0.29 |
| RF-score<sup>*, ◊, ∇</sup> | 0.62 | 2.00 | 0.05 |
| NN-score<sup>*, ◊, ∇</sup> | 0.58 | 1.38 | 0.03 |
| Ligand baseline<sup>*, ∇</sup> | 0.66 | 1.31 | 0.04 |

G, PDBbind general set; R, PDBbind refined set; \*, atom-level protein (pocket) representation; †, surface-level protein (pocket) representation; ◦, sequence-level protein representation; ∇, machine learning; ◊, docking.

**Table 4:** Statistical analysis for DUD-E virtual screening benchmark

|  |  | AUROC | EF<sub>1</sub> | BEDROC<sub>80.5</sub> |
| --- | --- | --- | --- | --- |
| DENVIS-G | DENVIS-R | *** | *** | *** |
| DENVIS-R | GNINA | n.s. | n.s. | n.s. |
| DENVIS-G | Gold | n.s. | *** | *** |
| DENVIS-G | Glide | ** | *** | *** |
| DENVIS-R | DeepDTA | *** | *** | *** |
| DENVIS-G | Ligand baseline | *** | *** | *** |
| DENVIS-R | Ligand baseline | *** | *** | *** |
| GNINA | Ligand baseline | *** | *** | *** |
| DeepDTA | Ligand baseline | ** | n.s. | n.s. |
| Vina | Ligand baseline | *** | *** | *** |
| RF-score | Ligand baseline | n.s. | n.s. | n.s. |
| NN-score | Ligand baseline | * | n.s. | n.s. |
| Gold | Ligand baseline | *** | *** | *** |
| Surflex | Ligand baseline | *** | *** | *** |
| Flexx | Ligand baseline | *** | *** | *** |
| Glide | Ligand baseline | *** | *** | *** |

\*, $p < 0.05$ ; \*\*, $p < 0.01$ ; \*\*\*, $p < 0.001$ ; n.s., non-significant. Abbreviations as in Table 3.

**Figure 3:**
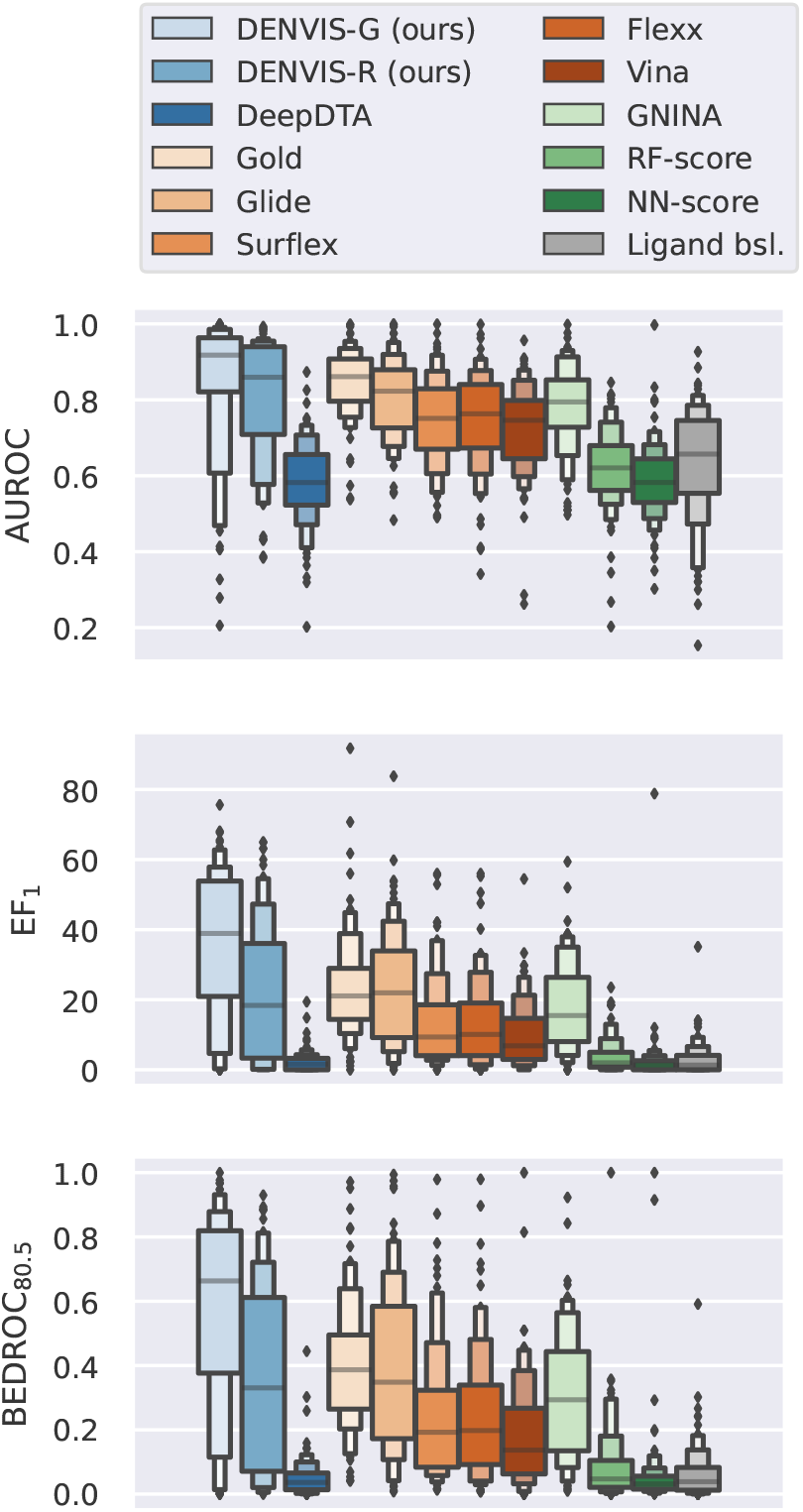
DUD-E virtual screening benchmark. The performance of several methods is shown using the AUROC, EF_1_, and BEDROC_80.5_ metrics. Box plots show the score distributions for the 102 DUD-E targets. Grey horizontal lines correspond to medians. Diamonds indicate outliers. Colour code: blues, machine learning-based; oranges, docking-based; greens, hybrid docking/machine learning-based; grey, ligand baseline.

DENVIS-G achieves the highest median AUROC performance followed by DENVIS-R, Gold, Glide, and GNINA. It also achieves the highest median EF_1_ and BEDROC_80.5_ scores. DENVIS-G significantly outperforms DENVIS-R in all three metrics. DENVIS-R achieves slightly higher median scores than GNINA, however the differences are below the significance level. It should be noted that both models have been trained on the same dataset (i.e., PDBbind refined set v. 2019).

Among the four docking algorithms, Gold and Glide achieve the highest performance. Both models are significantly outperformed by DENVIS-G, except for the AUROC metric which is comparable between DENVIS-G and Gold.

All models achieve a significantly higher AUROC score than our ligand-baseline, with RF-score being the only exception. With respect to EF_1_ and BEDROC_80.5_ scores, the following models do not significantly outperform the ligand-based baseline: RF-score, NN-score, and DeepDTA. For the DeepDTA algorithm, we additionally experiment with a model pre-trained on the DAVIS dataset, as was originally proposed by the authors^27^. This model performs worse than the one trained on the PDBbind refined set and considerably worse than our ligand-based baseline, achieving chance-level median AUROC, EF_1_ and BEDROC_80.5_ scores of 0.50, 0.0 and 0.0, respectively (not shown).

Figure 4 shows one-to-one scatter plot comparisons between (a) DENVIS-R and GNINA and (b) DENVIS-G and Gold for the three evaluation metrics of interest. Each point in these graphs corresponds to a single target in DUD-E. Across the 102 DUD-E targets, DENVIS-R achieves higher EF_1_ score than GNINA on 57 targets, and DENVIS-G achieves higher EF_1_ score than Gold on 73 targets. Extended per-target performance scores obtained with all algorithms in the benchmark are provided in the Supporting Information (Tables S1, S2, and S3 in the SI).

**Figure 4:**
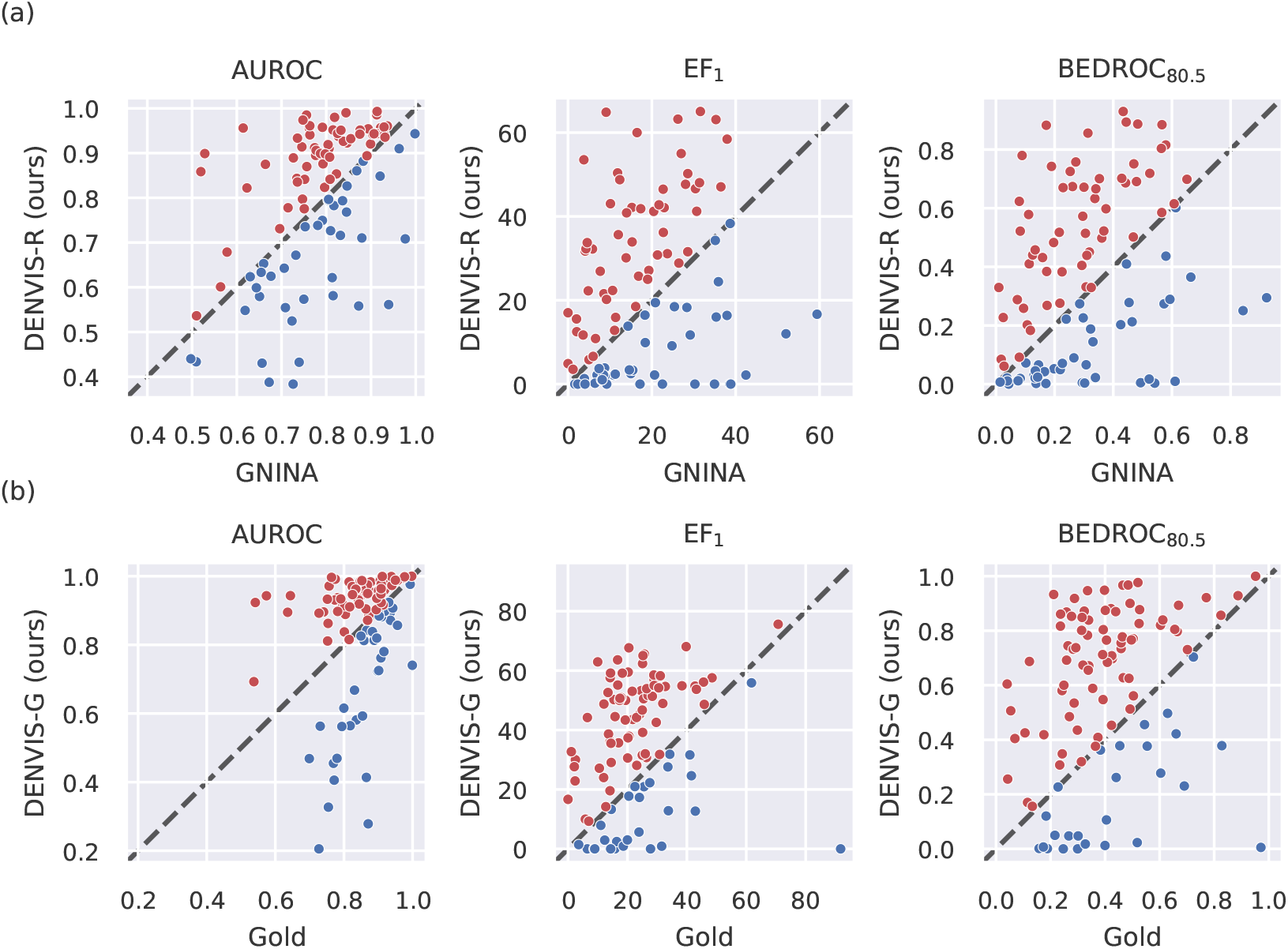
DUD-E per-target comparisons using AUROC, EF_1_ and BEDROC_80.5_ metrics. (a) Comparison between DENVIS-R and GNINA models. Both models are trained on the PDBbind refined set. (b) Comparison between DENVIS-G and Gold models. DENVIS-G is trained on the PDBbind general set, while Gold is a purely docking-based algorithms and therefore requires no training. Each point in the scatter plots corresponds to a single DUD-E target (*n* = 102), and is coloured to indicate the model that achieves the higher performance (DENVIS, red; GNINA/Gold, blue).

### Virtual screening performance benchmark: LIT-PCBA

We now evaluate the performance of our models on the LIT-PCBA benchmark^47^. This dataset contains multiple template files (i.e., pdb structures) for 13 out of 15 targets. In order to obtain a single prediction score when multiple templates are available per target, we predict the binding affinity of all template-ligand pairs and consider the template for which we predict the maximum binding affinity score^50^.

LIT-PCBA is a recently published benchmark and, hence, not many results have been yet made publicly available. Thus, we include in this analysis results with the two versions of DENVIS, as well as with DeepDTA^27^ and GNINA^21^, for which prediction scores have been published recently^50^. We also compare the performance of the three methods against our ligand baseline model. The results are shown in Figure 5 and median scores are also summarised in Table 5. We find that the two DENVIS models outperform DeepDTA but slightly underperform GNINA. The DENVIS and GNINA models achieve higher than random-level median EF_1_ performance, but this is not the case for the DeepDTA model. One-to-one comparisons between DENVIS and GNINA models are shown in Figure 6. Our DENVIS-R model achieves higher EF_1_ score than GNINA in six targets, GNINA achieves higher score in eight targets, and for one target the two models achieve equal scores. On the other hand, DENVIS-G achieves higher EF_1_ score than GNINA in eight targets and GNINA achieves higher score in seven targets. Extended per-target scores for all five models are provided in Tables S4, S5, and S6 in the SI. We do not perform statistical analyses for this dataset due to the small number of targets (see Methods section).

**Table 5:**
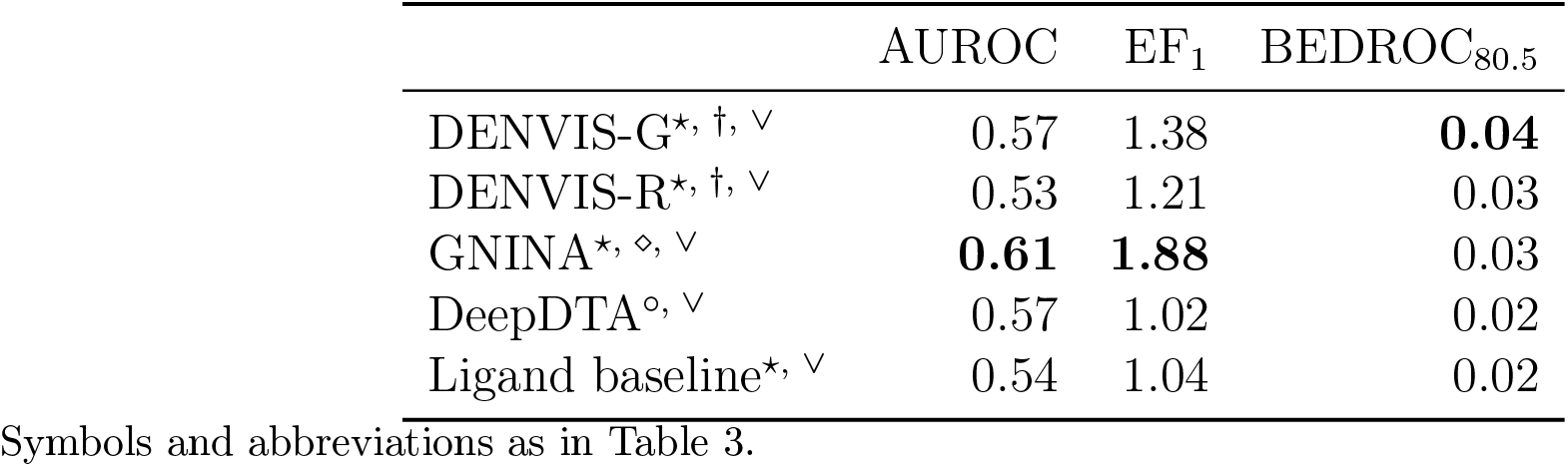
LIT-PCBA virtual screening benchmark summary (median scores)

**Figure 5:**
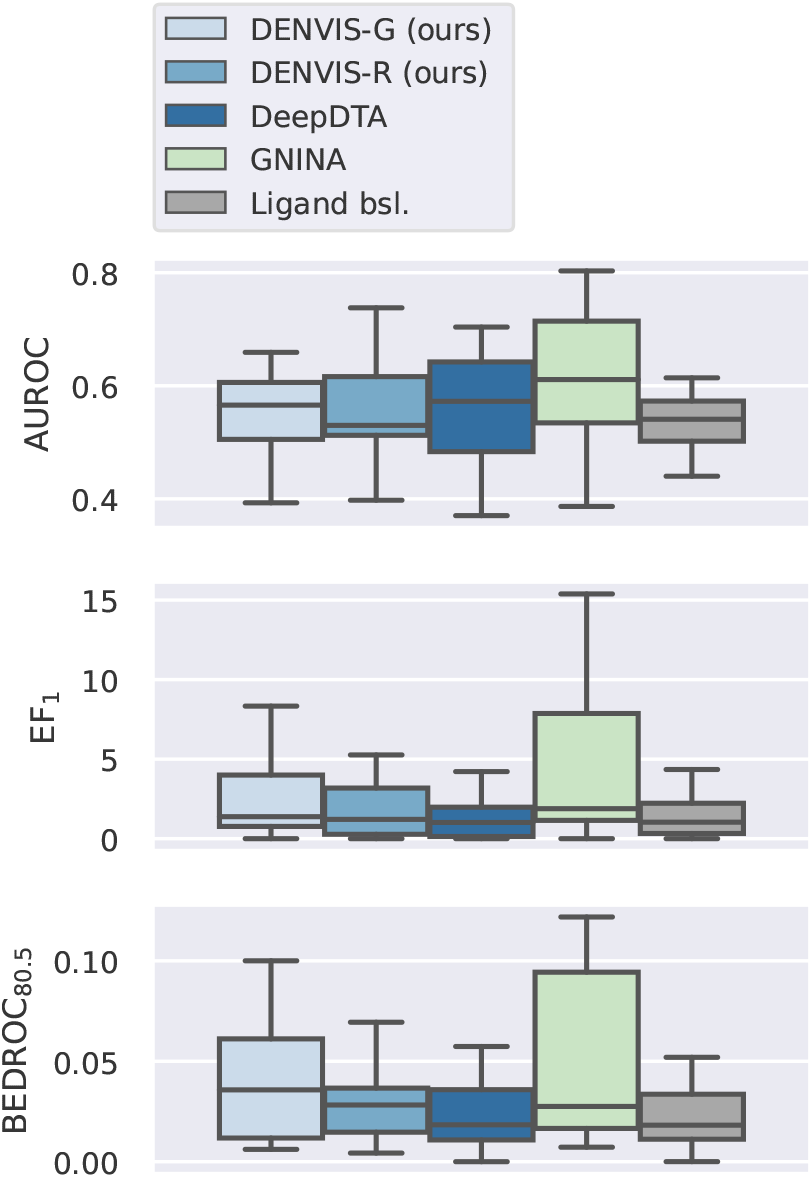
LIT-PCBA virtual screening benchmark. Box plots show score distributions for the 15 LIT-PCBA targets. Black horizontal lines correspond to medians.

**Figure 6:**
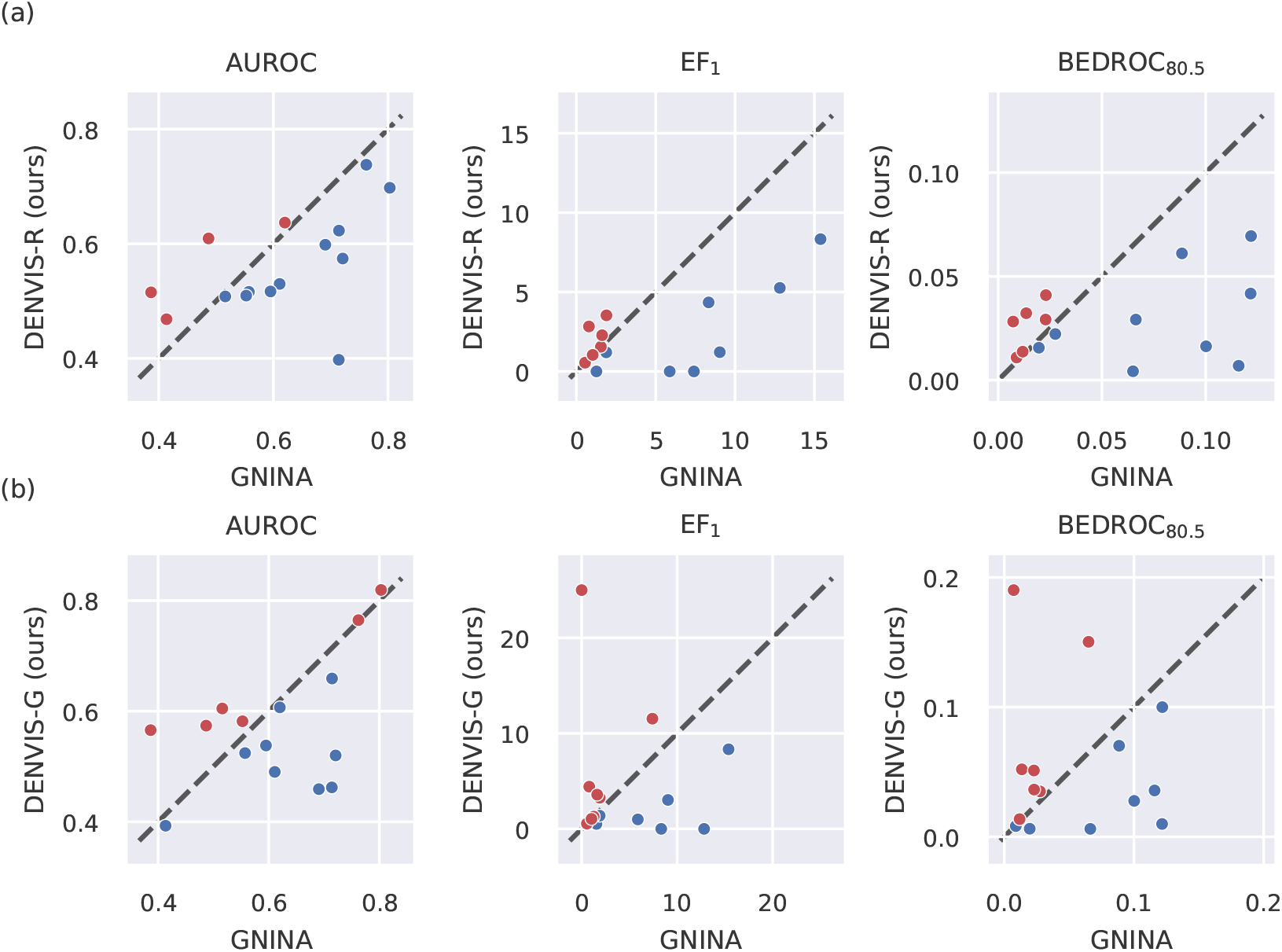
LIT-PCBA per-target comparisons using AUROC, EF_1_ and BEDROC_80.5_ metrics. (a) Comparison between DENVIS-R and GNINA models. (b) Comparison between DENVIS-G and GNINA models. GNINA is trained on the PDBbind refined set. Each point in the scatter plots corresponds to a single LIT-PCBA target (*n* = 15), and is coloured to indicate the model that achieves the higher performance (DENVIS, red; GNINA, blue).

### Virtual screening times benchmark

We now turn our attention to virtual screening (i.e., inference) times exhibited by a subset of algorithms in our benchmark: DENVIS, DeepDTA, Gold, FlexX, Surflex, Glide, and GNINA. For our method, we employ two strategies for screening, namely, naive and efficient screening. In the naive case, we compute the vector representations for every new protein pocket-ligand pair afresh, whereas with the efficient strategy we pre-compute the vector representations for each target protein pocket and store them to memory (see Methods section). In both cases, once the protein pocket and ligand vector representations have been (pre-)computed, they are fed to the interaction and final prediction layers to obtain the binding affinity prediction scores.

Average screening times for a protein pocket-ligand pair are presented in Table 6. For our method and DeepDTA we measure screening times on the DUD-E dataset using a Linux-operated machine (AMD Ryzen Threadripper 2970WX 24-Core Processor, 128 GB RAM, Ubuntu 18.04.4 LTS) and a single GPU (NVIDIA^®^ GeForce^®^ RTX 2080 Ti, 11 GB). We repeat all inference experiments 20 times and report average screening times (mean ± std). For Gold, FlexX, Surflex, Glide, and GNINA, we reproduce the times reported in the original references^21,61–64^. For Glide, we report the average time of the fastest mode (i.e., HTVS)^62^.

**Table 6:** Average virtual screening times per protein pocket-ligand pair

| Model | Implementation | Single prediction (ms) | Ensemble prediction (ms) |
| --- | --- | --- | --- |
| DENVIS (ours) | efficient | $0.314 \pm 0.008$ | $1.570 \pm 0.039$ |
| DENVIS (ours) | naive | $1.908 \pm 0.003$ | $9.540 \pm 0.014$ |
| DeepDTA | naive | $0.582 \pm 0.006$ | - |
| Gold<sup>61</sup> | - | $1.2 \times 10^3$ | - |
| FlexX<sup>63</sup> | - | $\sim 1 \times 10^3$ | - |
| Surflex<sup>64</sup> | - | $\sim 1\text{-}3 \times 10^3$ | - |
| Glide<sup>62</sup> | - | $\sim 0.5 \times 10^3$ | - |
| GNINA<sup>21</sup> | - | $2.6\text{ - }2.65 \times 10^4$ | $2.7 \times 10^4$ |

Using the efficient implementation our method exhibits an average screening time of 0.314 ms per protein pocket-ligand pair, which includes inference with one atom- and one surface-level model. A single DeepDTA model has nearly double inference times with an average of 0.582 ms. On the other hand, docking algorithms (i.e., Gold, FlexX, Surflex, Glide) exhibit three orders of magnitude higher screening times, that is, between 0.5 and 1.5s. Inference times with GNINA are four orders of magnitude higher than with our models, in the range of 26-27 s. The inference time of DENVIS scales linearly with the number of model instances in the ensemble, leading to an average of 1.570 ms per protein pocket-ligand pair when using ensembles comprising five atom- and surface-level models (in total 10 models). This exceeds the screening time of a single DeepDTA model by a factor of 3 approximately, while still being several orders of magnitude faster than docking algorithms and GNINA.

### Ablation studies

We now perform a series of ablation studies on the DUD-E dataset to assess the effect of several important design parameters of DENVIS on virtual screening performance. For the purposes of this analysis our baseline is the ensemble atom/surface-level model, for which both models in the ensemble comprise five model instances (i.e., multiple runs ensembling, Figure 1) trained on the PDBbind refined set. The GNN components are pre-trained using the M1 dataset and the metric learning approach described in the Methods section. Finally, during model training, we generate artificial negative examples as described in the Methods section. We refer to this model as *multi-ensemble (baseline)* in Figure 7, and *ME* in Tables 7 and 8.

**Table 7:** Ablation studies summary (DUD-E median scores)

|  | AUROC | EF<sub>1</sub> | BEDROC<sub>80.5</sub> |
| --- | --- | --- | --- |
| Multi-ensemble (baseline) | 0.86 | 18.37 | 0.33 |
| Single model prediction | 0.83 | 12.74 | 0.26 |
| Without pre-training | 0.87 | 16.67 | 0.34 |
| Without negative sampling | 0.63 | 1.96 | 0.05 |
| Binding classification | 0.78 | 9.39 | 0.20 |
| Level ensemble* | 0.87 | 18.48 | 0.35 |
| Atom-level* | 0.85 | 13.33 | 0.27 |
| Surface-level* | 0.78 | 11.07 | 0.23 |

\* Using subset of target protein pockets for which both atom-level and surface-level features are extracted (94/102 targets).

**Table 8:** Ablation studies statistical analysis (DUD-E dataset)

|  |  | AUROC | EF<sub>1</sub> | BEDROC<sub>80.5</sub> |
| --- | --- | --- | --- | --- |
| Multi-ensemble (baseline) | Single model prediction | *** | *** | *** |
|  | Without pre-training | n.s. | n.s. | n.s. |
|  | Without negative sampling | *** | *** | *** |
|  | Binding classification | *** | *** | *** |
| Level ensemble | Atom-level | n.s. | *** | *** |
|  | Surface-level | *** | *** | *** |

**Figure 7:**
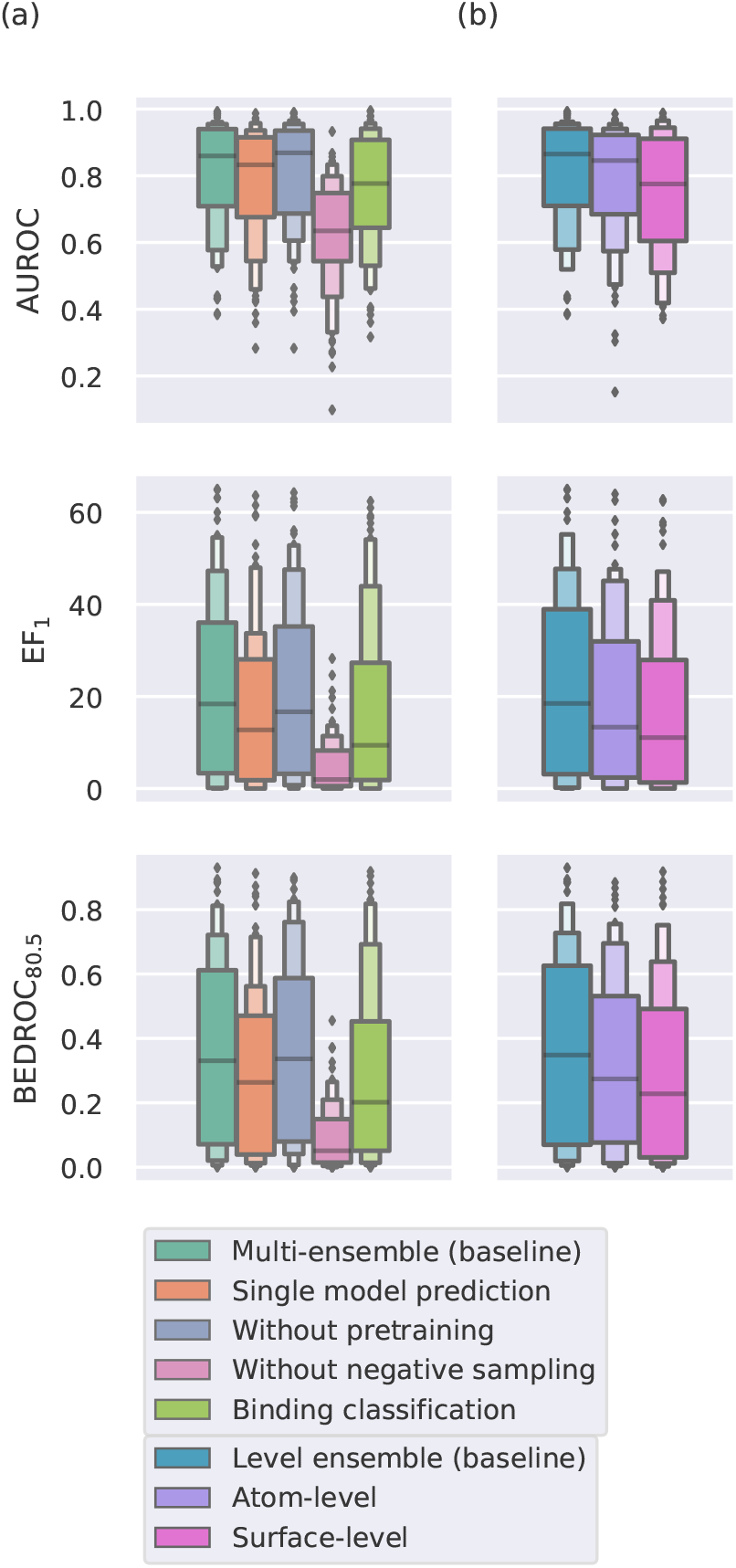
Ablation analysis (DUD-E dataset). (a) Multi-ensemble (baseline) refers to our complete method, the other models either have a component deactivated or some other modification (refer to main text for details). Box plots show score distributions for all DUD-E targets (*n* = 102). (b) Atom-level, surface-level and level ensemble comparison. Only targets for which protein surface preprocessing is successful are included (*n* = 94). All models are trained on PDBbind refined set for 600 epochs.

We experiment with the following modifications to our baseline model: 1) *single model prediction*, whereby the atom- and surface-level models comprise a single model instance (as opposed to five in the baseline model); 2) *without pre-training*, whereby all models are trained from scratch; 3) *without negative sampling*, whereby models are trained using positive examples only; and 4) *binding classification*, whereby our networks are trained with a binary cross-entropy loss using binary target labels indicating whether a ligand binds to the target protein pocket. Moreover, we compare the performance of *atom-level* (*AL*), *surface-level* (*SL*) and *level ensemble* (*LE*) models. We perform this analysis separately, since in this case we only include in the comparisons the target proteins for which surface-level preprocessing is successful (94/102 DUD-E targets, see Methods section).

The overall results of our ablation studies are presented in Figure 7 and median scores are also summarised in Table 7. The results from the statistical analysis are shown in Table 8. We observe that negative sampling is a key component of our method, since removing it leads to a dramatic decrease in performance. Using single model predictions (one for each of the atom- and surface-level models) also leads to a significant drop in performance. On the other hand, model pre-training does not seem to affect performance significantly (Table 8). Finally, the use of atom/surface-level model ensembles achieves significantly better performance than its atom-level and surface-level counterparts (Table 8).

We now perform further analysis to gain more insight into how combining multiple levels of protein pocket representation enhances the performance of our method. The precision-recall curves for the three types of models, namely, atom-level, surface-level and level ensemble, are shown in Figure 8(a). The curves correspond to aggregated targets for each of the three cases. It can be observed that the level ensemble model has better precision than its single-level counterparts, especially in the region of early ligand rankings (i.e., top-ranked ligands), which corresponds to the left-most area of the graph. Figure 8(b) shows an example of the scores predicted by the atom- and surface-level models for one target (Angiotensin-converting enzyme (ACE), PDB code: 3bkl). Each point in the scatter plot corresponds to one ligand and the colour code indicates whether the ligand is an active compound or a decoy. We observe that true binders tend to receive high scores by both models, which illustrates how the combination of the two models leads to higher precision. A similar behaviour is observed for the majority of target proteins in DUD-E. For the shown target, the atom-level, surface-level and level ensemble EF_1_ scores are 29.43, 11.35, and 32.27, respectively.

**Figure 8:**
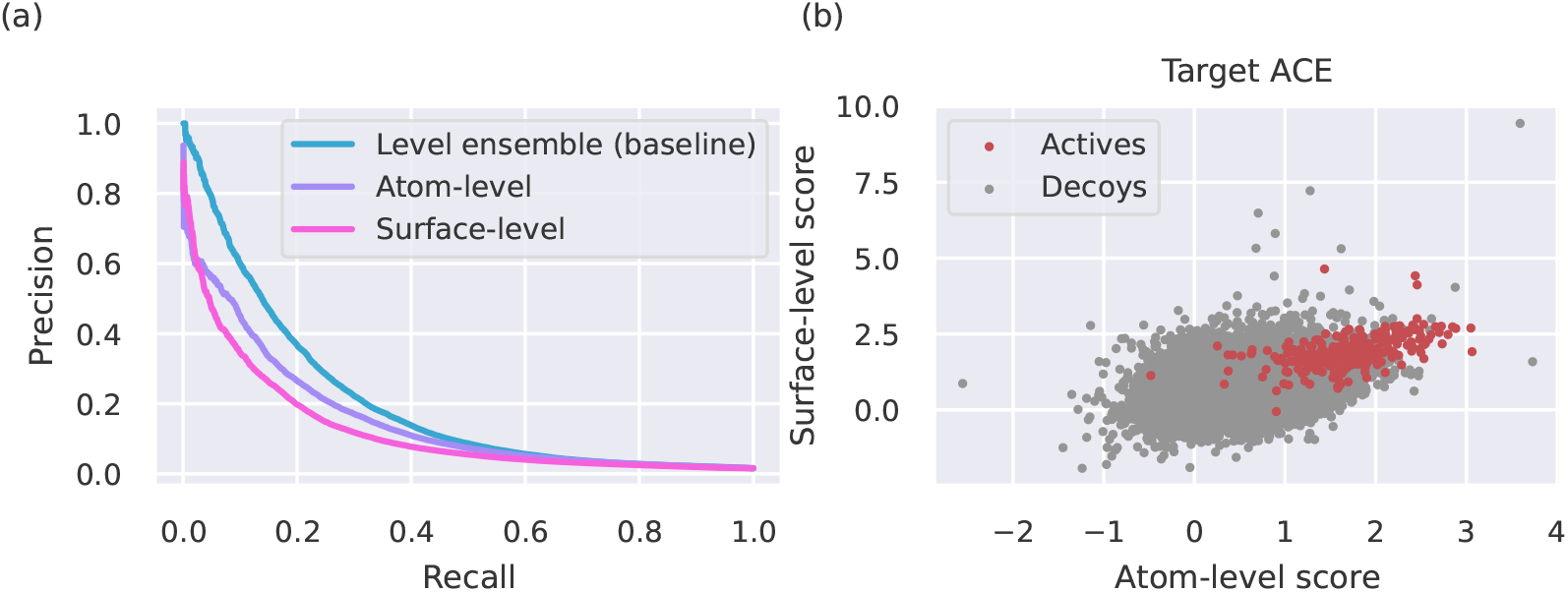
Combination of atom- and surface-level protein pocket representations (DUD-E dataset). (a) The precision-recall curves are shown for the three types of models: atom-level, surface-level, and level ensemble (baseline). Results from aggregated targets are shown (*n* = 94). (b) Scatter plot of predicted scores with atom- and surface-level models for Angiotensin-converting enzyme (ACE) target (PDB code: 3bkl). Each point corresponds to one ligand and the colour code indicates whether the ligand is an active compound or a decoy.

### Virtual screening evaluation metrics analysis

We observe that our three evaluation metrics, namely AUROC, EF_1_, and BEDROC_80.5_ are not always in perfect alignment in their assessments. For instance, on the DUD-E dataset, DeepDTA and NN-score achieve significantly higher AUROC scores than the ligand-based baseline, however they do not significantly outperform the baseline with respect to the EF_1_ and BEDROC_80.5_ metrics (Table 4).

To better understand the potential discrepancies between these three metrics, we study their relation and present the results of this analysis in Figure 9(a),(b). The two scatter plots correspond to the DENVIS-R model and show the linear relationship between (a) AUROC and EF_1_ and (b) AUROC and BEDROC_80.5_ scores, for the 102 targets in DUD-E. That is, each point in the scatter plots corresponds to one target protein in the database. In both cases, there is a significant positive correlation between the evaluation metrics, however it is not very strong (i.e., *r <* 0.80 in both cases). Moreover, for both models we observe outliers, which correspond to high EF_1_ and BEDROC_80.5_ scores (top-right corners) that lie far from the regression lines. This observation implies that the relationship of the metrics may in fact be non-linear.

**Figure 9:**
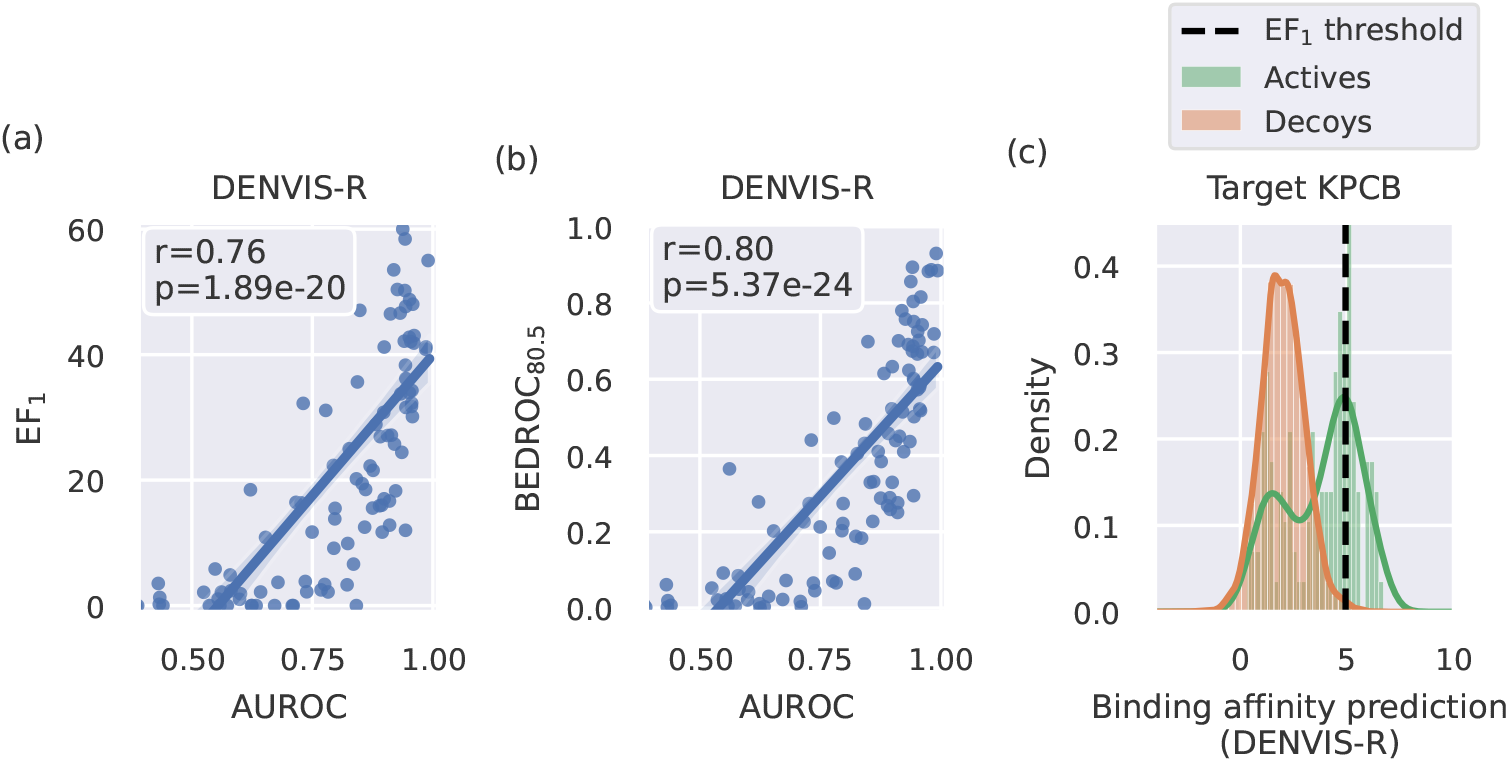
Virtual screening evaluation metric analysis (DUD-E dataset). (a)-(b) Linear relationship between (a) AUROC-EF_1_ and (b) AUROC-BEDROC_80.5_ scores using DENVIS-R model predictions. Linear regression fits and 95% confidence intervals are shown using bootstrapping (1000 iterations). Insets indicate the corresponding correlation coefficients (*r*) and significance values (*p*). (c) Example distribution of prediction scores using DENVIS-R for Protein kinase C beta (KPCB) target (PDB code: 2i0e). Distribution scores for the active and decoy sets are shown separately. The black vertical line denotes the cutoff threshold used by the EF_1_ score (i.e., top 1% of ranked ligands).

We attribute the relative discrepancies between the AUROC and EF_1_ metrics to the fact that they consider different (sub-)sets of the data: AUROC is a global metric which considers all available ligands for a given target, whereas EF_1_ only considers the top 1% of ranked ligands. On the other hand, the BEDROC is a global metric, that is, it considers all ligands, but uses exponential weighting to assign larger values to early-ranked ligands^57^. To illustrate how these different approaches can influence the measured performance, we provide an example for one target (Protein kinase C beta (KPCB), PDB code: 2i0e) in Figure 9(c) showing the distribution of binding affinity scores predicted by our model. Distributions of the active and decoy sets for this target are shown separately. We observe that, for the shown example, the two distributions overlap highly in the lower score range, however they are clearly separated above the EF_1_ cutoff threshold (black horizontal line). This is due to the active set having a heavy tail above this threshold. The resulting AUROC, EF_1_ and BEDROC_80.5_ scores for this example are 0.78 and 31.1, and 0.50, respectively.

## Discussion

### Virtual screening benchmarks

In this work, we introduce DENVIS, a GNN-based, end-to-end pipeline for drug virtual screening. In contrast with the majority of SBVS approaches, our method does not rely on docking simulations to estimate target-compound binding poses. We validate the performance of our model and compare it to competitive methods using two benchmark datasets. In DUD-E, we show that our approach achieves competitive performance to several commercial docking algorithms (i.e., Gold,^9^ Glide,^11^ Surflex,^10^ and Flexx^12^), when it is trained on the PDBbind refined set. Additionally, it achieves competitive performance to GNINA, a state-of-the-art research algorithm combining docking and a 3D CNN^21^. When trained on the PDBbind general set, our method outperforms all other methods in the benchmark by a large margin (Figure 3 and Tables 3, 4). On the LIT-PCBA benchmark, our DENVIS-R model achieves slightly inferior performance to GNINA (Figures 5, 6 and Table 5).

Even though the additional entries in the general set are generally considered to be of lower quality (i.e., resolution > 2.5 Å, complexes with covalent connections, complexes with multiple ligands in the binding pocket)^49^, our ensemble model trained on the PDBbind general set (DENVIS-G) outperforms its refined set counterpart (DENVIS-R) in both datasets. However, the performance difference is more profound in the case of DUD-E (Figures 3, 5 and Tables 3 4, 5). This observation is also in agreement with previous reports in the literature showing that the inclusion of more data in the training set leads to performance improvement for a binding affinity prediction network^20^. We attribute this finding to the fact that the additional entries in the PDBbind general set may cover a greater range of the protein and ligand input spaces and, thus, their inclusion in the training process may result in better generalisation of the models. In the ligand space, for instance, the general and refined sets contain approximately 11.3K and 3.1K unique ligands, respectively. It may also possible, however, that the improvement in performance may be partially attributed to a high similarity between the additional targets and/or ligands in the general set and those included in the DUD-E database^50^, hence, this aspect warrants further investigation in future work. With regards to computational requirements, the very fast inference time of our method renders it suitable for virtual screening of massive ligand libraries with minimal resources. By avoiding the intermediate docking step, our method exhibits three orders of magnitude faster screening times than state-of-the-art commercial docking algorithms, and four orders of magnitude faster screening times than GNINA (Table 6). We do not have access to screening times for AutoDock Vina, RF-score, and NN-score and therefore do not include them in the screening time benchmark. However, McNutt et al. ^21^ report an average docking time of 25 s with AutoDock Vina. Given that RF-score and NN-score build upon AutoDock Vina predictions, it is safe to assume that this figure is a lower bound for these two models. When compared to DeepDTA, a language-based method using whole-protein sequences and ligand string representations, which exhibits comparable inference times to our method, we show that we achieve dramatically better screening performance (Figures 3, 5 and Tables 3, 4, 5). Taking everything into consideration, it can be argued that our approach combines the best of two worlds, that is, the screening performance of state-of-the-art docking-based and hybrid docking/machine learning-based methods operating at the atomic level, and the throughput of purely machine learning-based models processing protein information at the sequence level. To further decrease the inference time of our method, we design an efficient screening strategy, whereby protein pocket and/or ligand vector representations are pre-computed and stored in memory. We show that this can further decrease the screening time by approximately a factor of 6. The ability of using such a type of efficient screening stems directly from the design of our algorithm, which does not take the protein-ligand complex as an input, but instead the two entities separately. This is not possible with docking-based methods, as in that case, the protein-ligand binding pose needs to be simulated separately for each protein-ligand pair.

### Novel aspects of DENVIS, comparison with related models and potential extensions

A few other methods have been recently proposed to circumvent the requirement for docking simulations. For example, Torng and Altman ^23^ and Li et al. ^34^ have introduced GNN architectures that model interactions between protein amino acids and small molecule atoms. One main difference between these two methods and ours is that we model interactions at the atomic level for target proteins, as opposed to amino acid sequence level. Another difference lies in the way that the protein and ligand vector representations are combined to produce the final binding affinity estimates. Torng and Altman ^23^ use a concatenation layer, whereas Li et al. ^34^ use a dual-attention mechanism in combination with an outer product layer. Attention-based mechanisms have also been proposed in the context of sequence-level models^31^.

It should be emphasised that our method is capable of producing accurate binding affinity estimates by considering interactions at the abstract space of protein pocket and compound vector representations rather than at the atomic level. This is due to the fact the protein-ligand interaction layer follows the global graph pooling operations in the two graphs. As a result, our method is binding pose invariant. Recently, a GNN-based method has been proposed for predicting ligand conformations for protein targets^68^. In the future, it shall be interesting to investigate the feasibility of combining this approach with our method in order to develop a system that can predict both binding pose and affinity simultaneously.

A crucial and novel aspect of our methodology is the use of negative data augmentation, whereby we generate artificial examples of non-binding protein pocket-ligand pairs during training. Our ablation analysis revealed that this is a key element of our approach with regards to its performance on a virtual screening task (Figure 7 and Table 8). This is somewhat expected, as the inclusion of negative examples forces our model to assign low scores to non-binding protein pocket-ligand pairs. In the opposite case, that is, when our model is only trained with binding protein-ligand complexes, the large majority of pocket-ligand pairs at test time (i.e., screening), which correspond to non-biding protein-ligand pairs, will be seen as out-of-distribution input data. For our negative sampling augmentation approach, we make the assumption that pockets and ligands from different complexes in the training set do not bind. In practice, this assumption may be violated, albeit with very small probability. It is nonetheless clear that the benefit of negative sampling on performance largely outweighs the potential negative effect of small label noise that may be introduced in the case of false negatives. Notably, we generate a different set of artificial negative examples in each training epoch, while keeping the positive/negative sample ratio fixed, thereby generating millions of negative samples during training. Although it is possible that other methods might benefit from the use of negative sampling augmentation, deploying this technique with docking-based methods would be likely infeasible, since in that case binding poses for all protein-ligand negative pairs would need to be estimated with docking simulations.

Another important contribution of our work is the representation of protein pocket information at two levels, namely, atomic and surface level. To our knowledge, such combination of protein pocket information at multiple levels of representation has not been previously reported. We show that this leads to a dramatic increase in performance (e.g., median EF_1_ improvement of 38.6% and 66.9% in DUD-E as compared to atom- and surface-level models, respectively). We observe that combining the two models leads to higher precision (Figure 8). Interestingly, this finding has been previously reported when combining different docking algorithms, a technique called consensus docking^69^. It is possible that combining fundamentally different models, such as different docking algorithms in consensus docking or, in our case, GNNs taking as inputs different types of protein pocket features, acts as a form of regularisation; errors made by one model can be counter-balanced by predictions from the other model. In our analysis, we observe that where both models agree, by assigning a high score to a specific ligand, the true probability of binding is usually high (Figure 8(b)). To the best of our knowledge, this is the first study combining multiple protein pocket representations for virtual drug screening. Here, we train separate models for the two types of representations and average their predictions at inference (i.e., late model fusion). An alternative approach could be to include both representations in a single multi-modal network and train it end-to-end.

Furthermore, in agreement with previous work^18,20,21,70^, we find that using model ensembles leads to an additional improvement in performance (average EF_1_ improvement of 29.0% in DUD-E). Since we initialise our model weights using network pre-training, the main difference between the base models in the ensemble is that they are trained using a different collection of artificially generated negative samples – and some stochasticity induced by data shuffling during training. One caveat of using model ensembles is that inference time increases linearly with the number of model instances in the ensemble. One possible way to overcome this limitation would be to use model ensembles in conjunction with knowledge distillation, whereby a student model could be trained to predict the output of a teacher ensemble model^71,72^.

Finally, we find that protein and ligand GNN pre-training using the TOUGH-M1 dataset and our metric learning-based approach does not yield a performance improvement, as one might have expected (Figure 7, Tables 7 and 8). Hu et al. ^39^ report an increase in performance using the MUV dataset^73^ and a combination of node-level and graph-level supervised pretraining. However, it should be noted that the datasets and validation strategies used in the two cases are different and, therefore, not directly comparable. The TOUGH-M1 dataset comprises more than 1M pairs of similar/dissimilar protein pockets and pairs of similar/dissimilar ligands. However, the number of unique proteins and ligands in the dataset is substantially lower, in the order of a few thousand (Table 1). It is possible that in order to achieve an improvement in virtual screening task performance with our metric learning-based pre-training approach, a much larger pre-training dataset is required. This would allow the protein pocket and ligand GNNs to have access to much richer information about the underlying protein and ligand structure distributions during the pre-training phase. Future research will further investigate alternative, self-supervised GNN pre-training strategies and their potential benefit to downstream virtual screening tasks.

### Dealing with biases in biochemical datasets

Machine learning-based methods for SBVS are often criticised for being prone to overfitting and failing to generalise to real-world scenarios^35–37,67,74–77^. Model validation is far from an easy task, as there exists strong evidence demonstrating that performance reported for machine learning-based algorithms is heavily affected by biases in the used biochemical datasets. A typical example is the comparative assessment of scoring functions (CASF) virtual screening benchmark^45,48,78^, which uses the refined and core subsets of the PDBbind database^49^ as the training and test sets, respectively. At the time it was first proposed^78^, the majority of scoring algorithms used empirical functions. High performance has since been reported with a wide variety of machine learning methods^48^, however there is striking evidence that such high performance may be largely attributed to the similarity between targets in the training and test sets^74,76^. Moreover, it has been found that binding affinity prediction algorithms trained on the PDBbind refined set might perform well on the core set but fail to generalise to other virtual screening datasets^75^. For all these reasons, an alternative benchmark is deemed necessary.

Other common forms of bias are analogue and decoy bias. Analogue bias refers to the case where a dataset may contain many analogue active ligands with the same chemotype. A machine learning model trained to recognise one such ligand as active for a given target is likely to assign high binding probability and/or affinity scores to the remaining ligands in the same scaffold^79^. On the other hand, decoy bias, or artificial enrichment, refers to the case where active and decoy sets are different in their basic molecular properties, including, but not limited to: molecular weight, LogP, number of hydrogen bond acceptors/donors, and number of rotatable bonds. A machine learning model trained on a partition of a dataset exhibiting decoy bias may trivially discriminate ligands based on their molecular properties and, thus, still perform well on an unseen partition of the dataset. Arguably, such a model may not learn any meaningful information about protein-ligand interaction that would allow it to generalise well to a different dataset or in a real-life application. To address this issue, the DUD-E database^46^ has been introduced, which uses carefully selected decoys with similar physicochemical properties to the active compounds. Yet, several reports show that both analogue and decoy biases are still present in DUD-E, and as a result simple baselines often perform competitively to sophisticated algorithms^37,75^. The high performance of such simple models may be attributed to the fact that the similarity between actives and decoys has been controlled at the level of single molecular descriptors. Yet, there may be systematic differences between actives and inactives in the second-order statistics of molecular properties (i.e., synergistic effect). Such synergistic differences may be exploited by simple baseline models considering only ligand properties, which can thus achieve artificially high performance^67^.

LIT-PCBA is a recently published dataset ^47^ that aims to address the decoy bias issues by employing an unbiasing technique that considers distances in the multidimensional molecular descriptor space^73^. Additionally, by containing single-assay data, all inactive compounds in the database have experimental support for inactivity. Another notable difference between DUD-E and LIT-PCBA is that the latter includes actives with typical potencies found in experimental screening decks. These are generally much smaller than the potencies of actives in DUD-E. In our experiments, we observe that all considered models, DENVIS, DeepDTA, and GNINA, achieve substantially lower performance in LIT-PCBA as compared to DUD-E. Yet, DENVIS and GNINA achieve better EF_1_ performance than our random baseline that only considers ligand information. It has been previously shown that the performance of both docking-based as well as hybrid docking/machine learning-based models is substantially lower in LIT-PCBA as compared to DUD-E^50^. This may be largely attributed to differences in active potency distributions between the two datasets, but also to other factors, such as incorrectly labelled actives in LIT-PCBA, and/or the decoy bias in DUD-E^37,50,67,75^.

In addition to unbiasing in the ligand space, an alternative approach to address the issue of dataset-specific biases, is to adopt a database splitting validation strategy^35^. With this approach, models are trained and tested on completely distinct databases^36,37^. The database splitting validation strategy^35^ is based on the assumption that analogue and decoy biases existing in a training dataset are unlikely to be shared with a test dataset, provided that the two sets have been constructed using different inclusion and filtering criteria. In our study, we follow this strategy and use the PDBbind database for training, and the DUD-E and LIT-PCBA databases for testing. An alternative, and perhaps more challenging strategy, is to cluster target proteins according to some similarity metric, for example, sequence or structural similarity, and evaluate performance on “unseen” targets^20,76^. The same procedure may be alternatively, or additionally, followed in the ligand space using a compound similarity metric, such as the Tanimoto coefficient^50^.

Last but not least, a common strategy to quantify the effect of dataset bias on performance is to develop appropriate baseline models. One such baseline concerns models that ignore the protein or ligand structure altogether^36,37^. To that end, we train a baseline model using ligand data only on the PDBbind refined set, which is the same dataset used to train all machine learning and hybrid docking/machine learning-based models in the benchmark. We observe that our ligand baseline achieves higher than chance-level performance (i.e., *AUROC >* 0.5 in both datasets). This finding verifies the hypothesis of across-dataset historical bias that may to some extent be transferred across datasets^50^. Interestingly, our ligand baseline performs competitively to many methods in our benchmark. Of note, DeepDTA, RF-score and NN-score show comparable performance to the baseline in terms of EF_1_ score (Figures 3, 5 and Tables 4, 5). This is somewhat surprising, especially when considering that DeepDTA is regarded as a state-of-the-art method for sequence-based virtual screening ^34,65,80^. Overall, our findings further reinforce the need for rigorous evaluation of machine learning models with appropriate baselines.

### Virtual screening evaluation metrics

As a final note, we observe a relative discrepancy between the three metrics that we use in our study, namely, AUROC and EF_1_ and BEDROC_80.5_ (Table 4). Although these three metrics seem to be positively correlated, their correlation is not very strong (i.e., *r* ≤ 0.80, Figure 9). From a statistical perspective, the EF measures recall (i.e., true positive rate), whereas AUROC and BEDROC consider both recall and fall-out (i.e., false positive rate). Furthermore, AUROC and BEDROC are global metrics, although the latter uses different weights for each ligand in the screening library based on its final ranking.

On the other hand, EF_1_ only considers the top 1% of ranked ligands for a given target (Figure 9). For virtual screening applications, it can be argued that the EF metric is more relevant, as it is desirable to experimentally validate only a fraction of the top-identified ligands for a given target^48^ and also put more emphasis on sensitivity/recall. Despite that, the EF metric suffers from its own limitations. Firstly, due to normalising by the total number of true binders for a given target (Equation 5), comparison across datasets, or even targets with the same dataset with different numbers of binders, may not be meaningful^4^. In such cases, the AUROC and BEDROC metric may be more appropriate. Alternatively, the metric can be normalised between in the range (0, 1).^50^. The normalisation with respect to number of true binders also gives rise to a second limitation. For targets with small numbers of actives, this metric can become heavily quantised. Nonetheless, both DUD-E and LIT-PCBA comprise a relatively large amount of active and decoy compounds for each target, and therefore the quantisation effect is minimal.

## Conclusion

In this work, we propose an end-to-end pipeline for high-throughput SBVS using ensemble GNNs and multiple levels of target protein pocket representation. By explicitly bypassing the requirement for docking simulations, our method achieves comparable performance to several state-of-the-art docking-based and hybrid docking/machine learning-based methods, yet with several orders of magnitude faster screening times. When compared to a purely machine learning-based sequence-level model with comparable inference times, our method achieves superior performance. In the future, we shall explore extensions of our model and further validate its performance with additional benchmark datasets.

## Data and software availability

We provide data and scripts required to reproduce all figures and tables in our manuscript and Supporting Information. Additionally, we provide support for using our methodology and trained models for virtual screening via a REST API: https://github.com/deeplab-ai/denvis.

We use the following publicly available datasets: TOUGH-M1, https://osf.io/6ngbs/; PDBbind v.2019, http://www.pdbbind-cn.org/; DUD-E, http://dude.docking.org/, and LIT-PCBA, https://drugdesign.unistra.fr/LIT-PCBA/.

Virtual screening results for DUD-E (GNINA. Vina, RF-score and NN-score) and LIT-PCBA (GNINA) are provided from the Koes research group at the Department of Computational and Systems Biology, University of Pittsburgh, and can be found in the following links: AutoDock Vina (DUD-E), http://bits.csb.pitt.edu/files/docked_dude.tar; GNINA (DUD-E), http://bits.csb.pitt.edu/files/defaultCNN_dude.tar.gz; GNINA (LIT-PCBA) http://bits.csb.pitt.edu/files/defaultCNN_litpcba.tar.gz; RF-score and NN-score (DUD-E), http://bits.csb.pitt.edu/files/rfnn_dude_scores.tgz.

## Acknowledgement

The authors would like to thank Liliane Mouawad from Curie Institute, Paris for kindly providing screening results for DUD-E with the four docking algorithms (i.e., Gold, Glide, Surflex and FlexX). We are also grateful to David Koes and Jocelyn Sunseri for kindly providing screening results for DUD-E with AutoDock Vina, GNINA, RF-score, and NN-score, and for LIT-PCBA with GNINA. We also thank Alexandros Pittis from the European Molecular Biology Laboratory (EMBL) for providing feedback on an earlier version of the manuscript. Finally, we thank NVIDIA Corporation for supporting this research by donating hardware via the “Applied Research Accelerator Program”. The authors received no specific funding for this work.

## Supporting Information

Empirical distribution of binding affinity metrics (*K*_*d*_, *K*_*i*_, and *IC*_50_) in PDBbind general set; model training dynamics; effect of number of base models in ensembles on performance; per-target performance scores (AUROC, EF_1_, and BEDROC_80.5_) for DUD-E and Lit-PCBA datasets. This material is available free of charge via the Internet at http://pubs.acs.org.

