## Supporting Information for "DENVIS: scalable and high-throughput virtual screening using graph neural networks with atomic and surface protein pocket features"

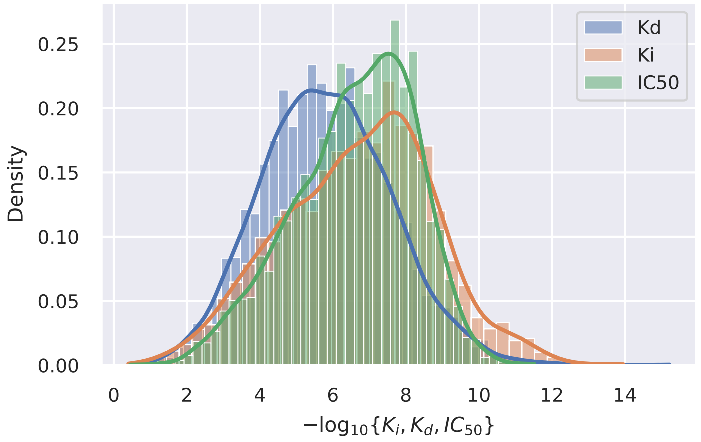


Figure S1: Empirical distributions of the three binding affinity metrics in PDBbind general set: $K_i$, $K_d$ and $IC_{50}$. Normalised histograms are shown along with kernel density estimates using Gaussian kernels.

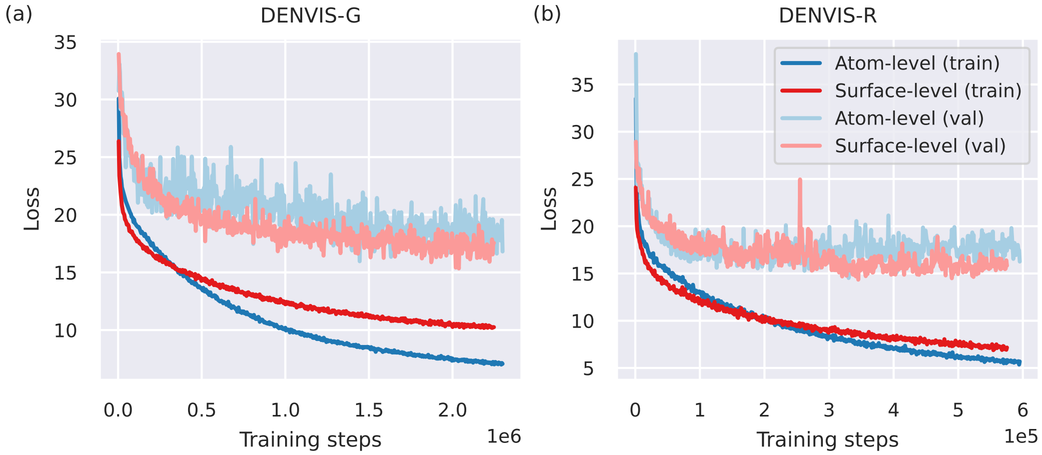


Figure S2: Model training dynamics. The evolution of training and validation losses is shown for the two types of models (atom- and surface-level). The shown dynamics correspond to a single model trained on PDBbind (a) general and (b) refined sets. Note that we do not perform early stopping during training. Instead, we select the checkpoint that yields the lowest validation loss after the training has completed.

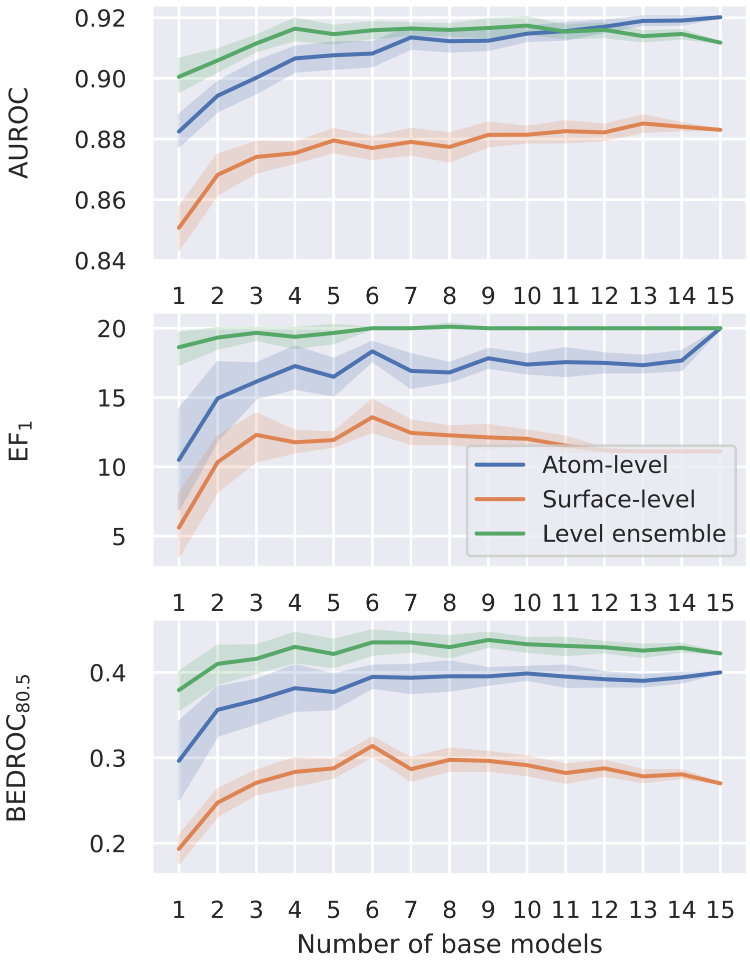


Figure S3: Performance against number of base models in ensembles. Performance is measured on the CASF-2016 benchmark (i.e., PDBBind core set v.2019). A total of 15 atom-level and surface-level models are independently trained on PDBbind refined set. To obtain confidence intervals, twenty independent runs are simulated. In each run, and for each number of base models in the horizontal axis, a subset of the 15 totally available models is sampled without replacement and the average (i.e., median) performance across targets is computed. Confidence intervals are not estimated for $n_{models} = 15$.

| | D-G | D-R | DDTA | Gold | Glide | Surflex | Flexx | Vina | GNINA | RF | NN | Bsl. |
|---|---|---|---|---|---|---|---|---|---|---|---|---|
| aa2ar | 0.84 | 0.43 | 0.59 | 0.87 | 0.80 | 0.77 | 0.74 | 0.65 | 0.74 | 0.50 | 0.54 | 0.57 |
| abl1 | 0.89 | 0.90 | 0.65 | 0.90 | 0.74 | 0.76 | 0.85 | 0.78 | 0.79 | 0.68 | 0.62 | 0.81 |
| ace | 0.90 | 0.96 | 0.41 | 0.74 | 0.72 | 0.77 | 0.61 | 0.56 | 0.61 | 0.67 | 0.75 | 0.40 |
| aces | 0.81 | 0.82 | 0.62 | 0.75 | 0.67 | 0.73 | 0.57 | 0.78 | 0.62 | 0.66 | 0.65 | 0.67 |
| ada | 0.99 | 0.99 | 0.61 | 0.91 | 0.68 | 0.83 | 0.55 | 0.57 | 0.76 | 0.61 | 0.58 | 0.61 |
| ada17 | 0.99 | 0.98 | 0.50 | 0.77 | 0.81 | 0.92 | 0.60 | 0.71 | 0.82 | 0.66 | 0.69 | 0.70 |
| adrb1 | 0.90 | 0.62 | 0.54 | 0.86 | 0.88 | 0.80 | 0.76 | 0.73 | 0.63 | 0.68 | 0.58 | 0.34 |
| adrb2 | 0.89 | 0.64 | 0.56 | 0.87 | 0.88 | 0.81 | 0.84 | 0.71 | 0.71 | 0.62 | 0.61 | 0.35 |
| akt1 | 0.88 | 0.96 | 0.79 | 0.91 | 0.78 | 0.63 | 0.73 | 0.76 | 0.76 | 0.63 | 0.49 | 0.71 |
| akt2 | 0.87 | 0.91 | 0.72 | 0.94 | 0.85 | 0.77 | 0.86 | 0.79 | 0.81 | 0.56 | 0.65 | 0.65 |
| aldr | 0.56 | 0.60 | 0.45 | 0.79 | 0.79 | 0.65 | 0.74 | 0.74 | 0.56 | 0.59 | 0.50 | 0.55 |
| ampc | 0.86 | 0.86 | 0.46 | 0.75 | 0.79 | 0.49 | 0.67 | 0.61 | 0.52 | 0.34 | 0.57 | 0.26 |
| andr | 0.94 | 0.88 | 0.57 | 0.57 | 0.84 | 0.57 | 0.41 | 0.63 | 0.79 | 0.64 | 0.57 | 0.69 |
| aofb | 0.58 | 0.54 | 0.47 | 0.84 | 0.76 | 0.68 | 0.71 | 0.78 | 0.51 | 0.53 | 0.56 | 0.54 |
| bace1 | 0.94 | 0.85 | 0.66 | 0.78 | 0.82 | 0.80 | 0.76 | 0.72 | 0.82 | 0.79 | 0.63 | 0.61 |
| braf | 0.96 | 0.95 | 0.67 | 0.91 | 0.86 | 0.70 | 0.89 | 0.86 | 0.83 | 0.63 | 0.56 | 0.80 |
| cah2 | 0.97 | 0.94 | 0.75 | 0.82 | 0.77 | 0.64 | 0.73 | 0.59 | 0.93 | 0.82 | 0.68 | 0.76 |
| casp3 | 0.92 | 0.63 | 0.53 | 0.91 | 0.76 | 0.70 | 0.70 | 0.70 | 0.65 | 0.54 | 0.63 | 0.51 |
| cdk2 | 0.96 | 0.90 | 0.66 | 0.85 | 0.88 | 0.64 | 0.76 | 0.72 | 0.80 | 0.55 | 0.59 | 0.77 |
| comt | 0.98 | 0.58 | 0.48 | 0.99 | 1.00 | 0.75 | 0.66 | 0.63 | 0.65 | 0.27 | 0.42 | 0.37 |
| cp2c9 | 0.56 | 0.55 | 0.57 | 0.82 | 0.64 | 0.59 | 0.70 | 0.62 | 0.62 | 0.59 | 0.53 | 0.58 |
| cp3a4 | 0.41 | 0.43 | 0.61 | 0.77 | 0.72 | 0.61 | 0.60 | 0.60 | 0.66 | 0.58 | 0.51 | 0.62 |
| csf1r | 0.97 | 0.91 | 0.74 | 0.87 | 0.80 | 0.61 | 0.84 | 0.69 | 0.78 | NaN | NaN | 0.80 |
| cxcr4 | 0.78 | 0.77 | 0.52 | 0.92 | 0.72 | 0.76 | 0.47 | 0.60 | 0.85 | 0.68 | 0.80 | 0.71 |
| def | 1.00 | 0.96 | 0.48 | 0.91 | 0.87 | 0.93 | 0.70 | 0.76 | 0.93 | 0.60 | 0.56 | 0.54 |
| dhi1 | 0.33 | 0.52 | 0.56 | 0.75 | 0.68 | 0.74 | 0.49 | 0.77 | 0.72 | 0.62 | 0.68 | 0.59 |
| dpp4 | 0.93 | 0.78 | 0.71 | 0.75 | 0.87 | 0.71 | 0.86 | 0.62 | 0.75 | 0.60 | 0.67 | 0.62 |
| drd3 | 0.56 | 0.55 | 0.55 | 0.73 | 0.72 | 0.77 | 0.67 | 0.75 | 0.71 | 0.66 | 0.60 | 0.66 |
| dyr | 0.96 | 0.94 | 0.61 | 0.92 | 0.87 | 0.87 | 0.67 | 0.77 | 0.90 | 0.55 | 0.68 | 0.67 |
| egfr | 0.93 | 0.91 | 0.58 | 0.84 | 0.84 | 0.67 | 0.83 | 0.64 | 0.77 | 0.54 | 0.55 | 0.69 |
| esr1 | 0.97 | 0.79 | 0.56 | 0.76 | 0.91 | 0.76 | 0.80 | 0.83 | 0.80 | 0.64 | 0.50 | 0.71 |
| esr2 | 0.93 | 0.75 | 0.55 | 0.79 | 0.94 | 0.76 | 0.84 | 0.80 | 0.79 | 0.65 | 0.50 | 0.71 |
| fa10 | 0.96 | 0.88 | 0.71 | 0.90 | 0.68 | 0.78 | 0.88 | 0.83 | 0.88 | 0.80 | 0.55 | 0.72 |
| fa7 | 0.99 | 0.99 | 0.67 | 0.97 | 0.91 | 0.98 | 0.97 | 0.91 | 0.91 | 0.85 | 0.70 | 0.55 |
| fabp4 | 0.95 | 0.90 | 0.53 | 0.87 | 0.85 | 0.76 | 0.87 | 0.78 | 0.53 | 0.66 | 0.30 | 0.44 |
| fak1 | 0.99 | 0.99 | 0.56 | 0.95 | 0.80 | 0.64 | 0.88 | 0.81 | 0.84 | 0.70 | 0.49 | 0.88 |
| fgfr1 | 0.47 | 0.56 | 0.53 | 0.70 | 0.48 | 0.58 | 0.59 | NaN | 0.94 | NaN | NaN | 0.49 |
| fkb1a | 0.95 | 0.89 | 0.40 | 0.84 | 0.87 | 0.80 | 0.58 | 0.77 | 0.78 | 0.65 | 0.57 | 0.47 |
| fnta | 0.90 | 0.74 | 0.58 | 0.78 | 0.75 | 0.69 | 0.76 | 0.65 | 0.78 | 0.74 | 0.57 | 0.73 |
| fpps | 0.74 | 0.71 | 0.20 | 1.00 | 0.55 | 0.52 | 0.82 | 0.29 | 0.98 | 0.20 | 1.00 | 0.32 |
| gcr | 0.69 | 0.80 | 0.64 | 0.54 | 0.72 | 0.67 | 0.54 | 0.64 | 0.75 | 0.60 | 0.64 | 0.75 |
| glcm | 0.72 | 0.68 | 0.48 | 0.90 | 0.65 | 0.82 | 0.69 | 0.49 | 0.58 | 0.49 | 0.60 | 0.49 |
| gria2 | 0.67 | 0.72 | 0.67 | 0.83 | 0.77 | 0.75 | 0.76 | 0.75 | 0.83 | 0.56 | 0.62 | 0.60 |
| grik1 | 0.90 | 0.84 | 0.60 | 0.92 | 0.90 | 0.81 | 0.87 | 0.59 | 0.73 | 0.46 | 0.68 | 0.50 |
| hdac2 | 0.96 | 0.92 | 0.61 | 0.88 | 0.63 | 0.85 | 0.81 | 0.85 | 0.80 | 0.81 | 0.54 | 0.75 |
| hdac8 | 0.98 | 0.94 | 0.60 | 0.87 | 0.88 | 0.84 | 0.76 | 0.82 | 0.83 | 0.75 | 0.53 | 0.77 |
| hivint | 0.47 | 0.44 | 0.44 | 0.78 | 0.72 | 0.67 | 0.67 | 0.71 | 0.50 | 0.48 | 0.47 | 0.52 |
| hivpr | 0.94 | 0.92 | 0.51 | 0.64 | 0.72 | 0.82 | 0.55 | 0.72 | 0.85 | 0.77 | 0.58 | 0.65 |
| hivrt | 0.84 | 0.58 | 0.45 | 0.80 | 0.83 | 0.67 | 0.61 | 0.68 | 0.82 | 0.50 | 0.56 | 0.56 |
| hmdh | 0.87 | 0.85 | 0.57 | 0.87 | 0.92 | 0.83 | 0.67 | 0.79 | 0.92 | 0.71 | 0.69 | 0.66 |
| hs90a | 0.99 | 0.89 | 0.57 | 0.85 | 0.65 | 0.49 | 0.34 | 0.26 | 0.73 | 0.39 | 0.44 | 0.52 |
| hxk4 | 0.84 | 0.78 | 0.54 | 0.88 | 0.57 | 0.56 | 0.67 | 0.57 | 0.82 | 0.62 | 0.63 | 0.74 |
| igf1r | 0.94 | 0.93 | 0.68 | 0.90 | 0.86 | 0.70 | 0.89 | 0.83 | 0.85 | 0.57 | 0.62 | 0.76 |
| inha | 0.86 | 0.62 | 0.38 | 0.96 | 0.56 | 0.84 | 0.69 | 0.72 | 0.68 | 0.66 | 0.54 | 0.57 |
| ital | 0.89 | 0.65 | 0.47 | 0.80 | 0.68 | 0.50 | 0.61 | 0.60 | 0.66 | 0.61 | 0.48 | 0.76 |
| jak2 | 0.99 | 0.94 | 0.70 | 0.94 | 0.88 | 0.73 | 0.83 | 0.78 | 0.91 | 0.51 | 0.63 | 0.84 |
| kif11 | 0.81 | 0.73 | 0.54 | 0.89 | 0.90 | 0.70 | 0.69 | 0.85 | 0.81 | 0.72 | 0.54 | 0.70 |
| kit | 0.96 | 0.96 | 0.67 | 0.83 | 0.65 | 0.60 | 0.79 | 0.78 | 0.79 | 0.65 | 0.56 | 0.74 |
| kith | 0.28 | 0.38 | 0.36 | 0.87 | 0.72 | 0.83 | 0.82 | 0.74 | 0.73 | 0.57 | 0.67 | 0.30 |
| kpcb | 0.82 | 0.78 | 0.59 | 0.90 | 0.86 | 0.84 | 0.80 | 0.77 | 0.71 | 0.69 | 0.49 | 0.60 |
| lck | 0.91 | 0.94 | 0.64 | 0.80 | 0.83 | 0.72 | 0.84 | 0.80 | 0.76 | 0.49 | 0.53 | 0.79 |
| lkha4 | 0.99 | 0.95 | 0.52 | 0.85 | 0.94 | 0.90 | 0.83 | 0.89 | 0.88 | 0.63 | 0.35 | 0.58 |
| mapk2 | 0.95 | 0.92 | 0.75 | 0.91 | 0.92 | 0.68 | 0.81 | 0.89 | 0.90 | 0.64 | 0.54 | 0.80 |
| mcr | 0.92 | 0.79 | 0.61 | 0.54 | 0.82 | 0.55 | 0.41 | 0.54 | 0.84 | 0.72 | 0.70 | 0.72 |
| met | 0.92 | 0.94 | 0.65 | 0.93 | 0.83 | 0.72 | 0.82 | 0.81 | 0.88 | 0.74 | 0.59 | 0.78 |
| mk01 | 0.97 | 0.96 | 0.65 | 0.94 | 0.84 | 0.74 | 0.87 | 0.86 | 0.76 | 0.68 | 0.61 | 0.83 |
| mk10 | 0.95 | 0.89 | 0.66 | 0.91 | 0.85 | 0.61 | 0.78 | 0.75 | 0.81 | 0.56 | 0.57 | 0.81 |
| mk14 | 0.98 | 0.91 | 0.71 | 0.82 | 0.72 | 0.55 | 0.85 | 0.74 | 0.75 | 0.64 | 0.51 | 0.81 |
| mmp13 | 1.00 | 0.97 | 0.57 | 0.76 | 0.77 | 0.83 | 0.63 | 0.66 | 0.75 | 0.66 | 0.63 | 0.84 |
| mp2k1 | 0.81 | 0.73 | 0.55 | 0.86 | 0.75 | 0.59 | 0.81 | 0.54 | 0.70 | 0.56 | 0.68 | 0.67 |
| nos1 | 0.41 | 0.60 | 0.47 | 0.87 | 0.75 | 0.72 | 0.76 | 0.59 | 0.64 | 0.54 | 0.60 | 0.45 |
| nram | 0.98 | 0.96 | 0.33 | 0.93 | 0.95 | 0.92 | 0.82 | 0.54 | 0.94 | 0.57 | 0.60 | 0.32 |
| pa2ga | 0.91 | 0.84 | 0.53 | 0.82 | 0.79 | 0.72 | 0.72 | 0.62 | 0.81 | 0.62 | 0.41 | 0.49 |
| parp1 | 0.92 | 0.86 | 0.64 | 0.80 | 0.95 | 0.79 | 0.84 | 0.86 | 0.87 | 0.58 | 0.62 | 0.66 |
| pde5a | 0.89 | 0.80 | 0.65 | 0.73 | 0.85 | 0.69 | 0.65 | 0.66 | 0.81 | 0.58 | 0.62 | 0.65 |
| pgh1 | 0.21 | 0.39 | 0.54 | 0.73 | 0.76 | 0.68 | 0.63 | 0.64 | 0.67 | 0.56 | 0.46 | 0.68 |
| pgh2 | 0.45 | 0.57 | 0.65 | 0.77 | 0.82 | 0.76 | 0.75 | 0.77 | 0.75 | 0.57 | 0.46 | 0.76 |
| plk1 | 0.93 | 0.94 | 0.60 | 0.90 | 0.91 | 0.62 | 0.81 | 0.64 | 0.91 | 0.60 | 0.47 | 0.63 |
| pnph | 1.00 | 0.99 | 0.83 | 0.94 | 0.76 | 0.68 | 0.74 | 0.88 | 0.91 | 0.52 | 0.59 | 0.76 |
| ppara | 0.95 | 0.87 | 0.50 | 0.85 | 0.73 | 0.86 | 0.75 | 0.86 | 0.76 | 0.62 | 0.54 | 0.57 |
| ppard | 0.96 | 0.93 | 0.52 | 0.84 | 0.66 | 0.85 | 0.68 | 0.76 | 0.74 | 0.54 | 0.51 | 0.58 |
| pparg | 0.92 | 0.84 | 0.51 | 0.84 | 0.74 | 0.79 | 0.67 | 0.79 | 0.74 | 0.71 | 0.53 | 0.61 |
| prgr | 0.90 | 0.82 | 0.62 | 0.64 | 0.84 | 0.63 | 0.68 | 0.67 | 0.80 | 0.67 | 0.65 | 0.79 |
| ptn1 | 0.76 | 0.62 | 0.58 | 0.91 | 0.84 | 0.85 | 0.82 | 0.83 | 0.81 | 0.62 | 0.60 | 0.65 |

| | | | | | | | | | | | | |
|---|---|---|---|---|---|---|---|---|---|---|---|---|
| pur2 | 1.00 | 0.94 | 0.87 | 1.00 | 0.99 | 1.00 | 1.00 | 0.91 | 1.00 | 0.74 | 0.83 | 0.75 |
| pygm | 0.62 | 0.43 | 0.42 | 0.80 | 0.56 | 0.55 | 0.74 | 0.60 | 0.51 | 0.62 | 0.67 | 0.55 |
| pyrd | 0.59 | 0.67 | 0.53 | 0.85 | 0.88 | 0.77 | 0.83 | 0.83 | 0.73 | 0.78 | 0.49 | 0.73 |
| reni | 0.83 | 0.74 | 0.62 | 0.85 | 0.84 | 0.90 | 0.73 | 0.66 | 0.75 | 0.71 | 0.65 | 0.48 |
| rock1 | 0.88 | 0.83 | 0.65 | 0.90 | 0.84 | 0.74 | 0.91 | 0.72 | 0.85 | 0.57 | 0.55 | 0.69 |
| rxra | 0.95 | 0.94 | 0.65 | 0.82 | 0.97 | 0.84 | 0.80 | 0.81 | 0.93 | 0.47 | 0.52 | 0.93 |
| sahh | 0.90 | 0.84 | 0.32 | 0.89 | 0.98 | 0.91 | 0.91 | 0.80 | 0.75 | 0.54 | 0.71 | 0.15 |
| src | 0.93 | 0.93 | 0.72 | 0.77 | 0.74 | 0.62 | 0.80 | 0.65 | 0.84 | 0.53 | 0.55 | 0.78 |
| tgfr1 | 0.99 | 0.94 | 0.71 | 0.96 | 0.96 | 0.77 | 0.89 | 0.90 | 0.93 | 0.58 | 0.49 | 0.57 |
| thb | 0.92 | 0.71 | 0.51 | 0.91 | 0.85 | 0.87 | 0.83 | 0.82 | 0.88 | 0.61 | 0.38 | 0.71 |
| thrb | 0.99 | 0.96 | 0.66 | 0.85 | 0.92 | 0.88 | 0.88 | 0.77 | 0.92 | 0.81 | 0.79 | 0.58 |
| try1 | 0.98 | 0.95 | 0.74 | 0.88 | 0.91 | 0.94 | 0.90 | 0.80 | 0.89 | 0.78 | 0.72 | 0.61 |
| tryb1 | 0.73 | 0.87 | 0.67 | 0.90 | 0.88 | 0.91 | 0.86 | 0.71 | 0.66 | 0.72 | 0.66 | 0.49 |
| tysy | 0.89 | 0.56 | 0.71 | 0.94 | 0.93 | 0.84 | 0.84 | 0.87 | 0.87 | 0.68 | 0.64 | 0.70 |
| urok | 0.98 | 0.95 | 0.72 | 0.91 | 0.92 | 0.76 | 0.93 | 0.77 | 0.82 | 0.67 | 0.60 | 0.69 |
| vgfr2 | 0.82 | 0.94 | 0.70 | 0.82 | 0.68 | 0.69 | 0.85 | 0.77 | 0.83 | 0.71 | 0.50 | 0.73 |
| wee1 | 1.00 | 0.91 | 0.49 | 0.98 | 1.00 | 0.95 | 0.96 | 0.96 | 0.96 | 0.77 | 0.59 | 0.78 |
| xiap | 0.91 | 0.89 | 0.58 | 0.94 | 0.82 | 0.90 | 0.91 | 0.73 | 0.89 | 0.74 | 0.80 | 0.41 |

Table S1: DUD-E per-target AUROC scores. Abbreviations: D-G, DENVIS-PDBbind general set; D-R, DENVIS-PDBbind refind set; DDTA, Deep DTA; RF, RF-score; NN, NN-score; Bsl., Ligand baseline.

| | D-G | D-R | DDTA | Gold | Glide | Surflex | Flexx | Vina | GNINA | RF | NN | Bsl. |
|---|---|---|---|---|---|---|---|---|---|---|---|---|
| aa2ar | 35.68 | 0.21 | 1.66 | 17.01 | 5.21 | 19.71 | 6.02 | 2.49 | 6.43 | 2.28 | 0.62 | 1.24 |
| abl1 | 51.65 | 41.21 | 8.24 | 26.92 | 17.42 | 17.03 | 27.78 | 13.26 | 20.44 | 11.05 | 2.21 | 4.40 |
| ace | 29.08 | 32.27 | 0.00 | 14.54 | 5.42 | 2.48 | 1.42 | 2.86 | 4.29 | 5.71 | 4.64 | 0.00 |
| aces | 20.97 | 3.31 | 8.83 | 22.52 | 8.13 | 20.31 | 1.35 | 14.57 | 15.45 | 3.97 | 1.55 | 3.31 |
| ada | 59.14 | 40.86 | 0.00 | 18.28 | 5.38 | 9.68 | 10.75 | 1.08 | 13.98 | 4.30 | 0.00 | 0.00 |
| ada17 | 67.67 | 65.04 | 0.56 | 20.49 | 28.91 | 35.90 | 1.89 | 28.01 | 31.58 | 2.82 | 3.76 | 2.07 |
| adrb1 | 22.27 | 0.00 | 2.43 | 27.53 | 18.70 | 15.38 | 9.35 | 4.05 | 4.05 | 0.40 | 2.02 | 0.40 |
| adrb2 | 30.74 | 2.16 | 2.16 | 26.41 | 27.71 | 20.35 | 20.00 | 3.46 | 6.93 | 0.87 | 3.46 | 1.30 |
| akt1 | 17.75 | 31.74 | 4.44 | 20.48 | 10.92 | 4.44 | 5.17 | 5.46 | 4.10 | 1.37 | 0.00 | 1.37 |
| akt2 | 31.62 | 12.82 | 3.42 | 41.03 | 24.35 | 14.53 | 13.68 | 26.50 | 11.11 | 0.85 | 1.71 | 4.27 |
| aldr | 5.66 | 1.89 | 0.00 | 23.90 | 27.67 | 7.55 | 11.95 | 9.43 | 8.81 | 1.26 | 0.00 | 1.89 |
| ampc | 16.67 | 12.50 | 0.00 | 0.00 | 2.08 | 0.00 | 0.00 | 0.00 | 2.08 | 0.00 | 0.00 | 0.00 |
| andr | 32.71 | 21.56 | 1.49 | 1.12 | 25.70 | 4.09 | 0.00 | 17.84 | 8.55 | 1.49 | 2.60 | 2.23 |
| aofb | 2.46 | 0.00 | 0.82 | 16.39 | 6.67 | 4.92 | 5.26 | 6.56 | 1.64 | 0.82 | 0.00 | 4.92 |
| bace1 | 50.53 | 19.43 | 0.35 | 25.44 | 14.13 | 17.31 | 11.11 | 4.59 | 20.85 | 18.37 | 2.12 | 0.00 |
| braf | 46.71 | 42.76 | 3.95 | 25.00 | 29.80 | 5.26 | 23.03 | 17.11 | 21.71 | 12.50 | 0.66 | 1.32 |
| cah2 | 63.62 | 63.21 | 1.42 | 16.70 | 3.89 | 3.05 | 7.39 | 0.00 | 26.22 | 18.50 | 3.66 | 14.02 |
| casp3 | 39.20 | 0.00 | 0.00 | 25.13 | 27.78 | 12.56 | 22.11 | 1.52 | 17.17 | 2.02 | 4.04 | 0.00 |
| cdk2 | 44.51 | 30.80 | 2.74 | 15.82 | 29.87 | 3.16 | 11.80 | 9.51 | 21.35 | 0.85 | 1.48 | 9.49 |
| comt | 75.61 | 4.88 | 0.00 | 70.73 | 83.78 | 2.44 | 9.76 | 4.88 | 0.00 | 0.00 | 4.88 | 0.00 |
| cp2c9 | 10.00 | 5.83 | 1.67 | 5.83 | 0.88 | 3.33 | 9.09 | 3.33 | 5.00 | 5.00 | 0.00 | 0.00 |
| cp3a4 | 2.94 | 3.53 | 3.53 | 12.35 | 11.80 | 8.82 | 4.19 | 1.80 | 1.20 | 3.59 | 0.00 | 2.35 |
| csf1r | 43.37 | 27.11 | 4.82 | 19.28 | 18.40 | 3.01 | 24.54 | 1.20 | 19.28 | NaN | NaN | 4.22 |
| cxcr4 | 30.00 | 2.50 | 2.50 | 2.50 | 0.00 | 15.00 | 0.00 | 0.00 | 15.00 | 0.00 | 5.00 | 0.00 |
| def | 53.92 | 48.04 | 0.00 | 26.47 | 5.13 | 32.35 | 0.00 | 9.80 | 31.37 | 4.90 | 1.96 | 0.00 |
| dhi1 | 0.00 | 2.12 | 1.21 | 8.79 | 7.03 | 3.64 | 0.63 | 4.24 | 10.91 | 6.36 | 1.82 | 1.21 |
| dpp4 | 30.58 | 3.38 | 4.50 | 20.08 | 11.84 | 6.38 | 11.84 | 0.56 | 14.63 | 0.75 | 3.19 | 1.88 |
| drd3 | 7.92 | 1.04 | 1.04 | 11.04 | 1.69 | 7.08 | 2.51 | 5.02 | 8.16 | 3.77 | 1.26 | 4.17 |
| dyr | 57.58 | 50.22 | 2.60 | 48.48 | 30.87 | 25.54 | 10.43 | 7.36 | 28.57 | 1.30 | 1.30 | 1.73 |
| egfr | 62.36 | 46.49 | 2.58 | 25.09 | 24.95 | 13.65 | 19.29 | 5.72 | 22.69 | 2.58 | 2.77 | 8.86 |
| esr1 | 53.26 | 9.14 | 2.35 | 24.61 | 48.78 | 27.15 | 31.73 | 18.54 | 24.80 | 0.00 | 1.31 | 0.52 |
| esr2 | 50.68 | 11.72 | 1.91 | 17.49 | 44.26 | 21.80 | 31.02 | 14.99 | 29.16 | 0.00 | 0.27 | 0.82 |
| fa10 | 31.84 | 28.86 | 4.47 | 34.26 | 13.74 | 8.57 | 33.08 | 15.83 | 26.44 | 13.04 | 0.93 | 2.42 |
| fa7 | 56.14 | 41.23 | 3.51 | 45.61 | 44.04 | 42.11 | 55.26 | 11.40 | 30.70 | 18.42 | 0.00 | 0.00 |
| fabp4 | 48.94 | 17.02 | 0.00 | 31.91 | 25.58 | 2.13 | 17.02 | 29.79 | 0.00 | 0.00 | 0.00 | 0.00 |
| fak1 | 55.00 | 55.00 | 2.00 | 29.00 | 20.00 | 5.00 | 15.00 | 19.00 | 27.00 | 2.00 | 0.00 | 3.00 |
| fgfr1 | 1.44 | 2.16 | 2.16 | 3.60 | 0.00 | 0.72 | 0.76 | 1.44 | 42.45 | 1.44 | 1.44 | 1.44 |
| fkb1a | 31.53 | 11.71 | 0.00 | 25.00 | 41.82 | 9.91 | 0.00 | 7.21 | 3.60 | 1.80 | 0.00 | 0.90 |
| fnta | 53.04 | 2.20 | 0.84 | 21.62 | 6.85 | 4.94 | 6.26 | 2.70 | 8.45 | 8.11 | 1.52 | 1.18 |
| fpps | 0.00 | 0.00 | 0.00 | 91.76 | 1.23 | 1.18 | 1.18 | 0.00 | 38.82 | 0.00 | 78.82 | 0.00 |
| gcr | 9.30 | 15.50 | 0.78 | 6.98 | 16.56 | 15.12 | 7.63 | 15.12 | 15.50 | 2.33 | 2.33 | 3.88 |
| glcm | 0.00 | 3.70 | 1.85 | 27.78 | 11.11 | 31.48 | 5.56 | 0.00 | 7.41 | 0.00 | 1.85 | 0.00 |
| gria2 | 13.29 | 16.46 | 1.90 | 14.56 | 33.33 | 8.23 | 7.59 | 10.76 | 18.35 | 4.43 | 8.86 | 2.53 |
| grik1 | 42.57 | 35.64 | 0.99 | 29.70 | 20.00 | 6.93 | 19.80 | 4.00 | 12.00 | 2.00 | 4.00 | 0.99 |
| hdac2 | 55.14 | 53.51 | 2.70 | 16.76 | 15.84 | 12.97 | 16.30 | 11.89 | 3.78 | 2.70 | 2.16 | 3.78 |
| hdac8 | 62.94 | 60.00 | 1.76 | 10.00 | 2.45 | 5.29 | 12.94 | 20.00 | 16.47 | 3.53 | 0.59 | 5.29 |
| hivint | 0.00 | 0.00 | 0.00 | 9.00 | 2.06 | 3.00 | 4.00 | 2.00 | 0.00 | 0.00 | 0.00 | 0.00 |
| hivpr | 50.93 | 18.28 | 1.49 | 17.91 | 9.14 | 5.04 | 2.08 | 4.10 | 28.36 | 23.51 | 0.19 | 7.09 |
| hivrt | 14.20 | 2.37 | 0.59 | 12.72 | 22.94 | 4.73 | 2.08 | 4.15 | 11.28 | 0.30 | 2.08 | 6.21 |
| hmdh | 45.29 | 47.06 | 0.00 | 24.12 | 39.41 | 17.06 | 1.76 | 4.12 | 36.47 | 4.71 | 1.76 | 0.00 |
| hs90a | 38.64 | 15.91 | 2.27 | 13.64 | 1.16 | 1.14 | 2.35 | 0.00 | 11.36 | 0.00 | 2.27 | 0.00 |
| hxk4 | 19.57 | 2.17 | 1.09 | 14.13 | 22.83 | 0.00 | 4.35 | 3.26 | 20.65 | 0.00 | 0.00 | 14.13 |
| igf1r | 51.35 | 46.62 | 3.38 | 28.38 | 32.65 | 10.81 | 27.70 | 16.22 | 30.41 | 12.84 | 2.70 | 1.35 |
| inha | 20.93 | 0.00 | 0.00 | 25.58 | 9.30 | 4.65 | 9.30 | 6.98 | 2.33 | 0.00 | 0.00 | 0.00 |
| ital | 44.20 | 10.87 | 0.00 | 6.52 | 8.91 | 2.90 | 1.45 | 0.00 | 6.57 | 0.73 | 0.73 | 2.90 |
| jak2 | 57.01 | 47.66 | 6.54 | 28.97 | 33.77 | 14.95 | 8.41 | 15.89 | 28.04 | 8.41 | 2.80 | 7.48 |
| kif11 | 27.59 | 16.38 | 0.00 | 33.62 | 38.79 | 6.03 | 2.59 | 24.14 | 37.93 | 14.66 | 0.00 | 4.31 |
| kit | 35.54 | 30.12 | 4.22 | 14.46 | 6.10 | 3.01 | 7.88 | 4.82 | 13.86 | 6.63 | 2.41 | 4.22 |
| kith | 0.00 | 0.00 | 0.00 | 15.79 | 30.77 | 42.11 | 0.00 | 33.33 | 38.60 | 1.75 | 0.00 | 0.00 |
| kpcb | 58.52 | 31.11 | 0.00 | 28.89 | 52.46 | 27.41 | 27.48 | 28.15 | 23.70 | 0.00 | 2.22 | 6.67 |
| lck | 50.24 | 42.14 | 3.81 | 15.48 | 25.12 | 13.57 | 18.12 | 10.00 | 15.24 | 3.57 | 0.71 | 2.14 |
| lkha4 | 52.63 | 33.92 | 0.00 | 13.45 | 28.24 | 18.71 | 7.60 | 14.71 | 15.29 | 0.00 | 0.00 | 1.17 |

| | | | | | | | | | | | | |
|---|---|---|---|---|---|---|---|---|---|---|---|---|
| mapk2 | 43.56 | 25.74 | 14.85 | 21.78 | 27.72 | 2.97 | 6.93 | 13.86 | 16.83 | 1.98 | 1.98 | 3.96 |
| mcr | 27.66 | 22.34 | 1.06 | 2.13 | 29.41 | 2.13 | 0.00 | 7.45 | 10.64 | 4.26 | 9.57 | 4.26 |
| met | 68.07 | 58.43 | 1.20 | 39.76 | 27.71 | 9.04 | 18.07 | 12.05 | 37.95 | 19.28 | 1.20 | 1.20 |
| mk01 | 59.49 | 43.04 | 1.27 | 20.25 | 12.66 | 1.27 | 21.52 | 3.80 | 10.13 | 5.06 | 1.27 | 1.27 |
| mk10 | 44.23 | 26.92 | 3.85 | 23.08 | 12.50 | 3.85 | 14.42 | 7.69 | 7.69 | 4.81 | 0.96 | 4.81 |
| mk14 | 57.61 | 27.16 | 2.42 | 14.19 | 22.15 | 3.29 | 14.19 | 6.57 | 19.38 | 7.44 | 0.52 | 4.84 |
| mmp13 | 65.56 | 64.86 | 1.57 | 25.70 | 18.41 | 12.76 | 4.20 | 4.02 | 9.09 | 5.59 | 2.10 | 9.27 |
| mp2k1 | 28.10 | 32.23 | 0.00 | 23.14 | 13.22 | 0.00 | 8.33 | 0.00 | 5.83 | 0.83 | 5.00 | 6.61 |
| nos1 | 3.00 | 1.00 | 1.00 | 20.00 | 14.29 | 24.00 | 7.07 | 2.00 | 8.00 | 0.00 | 2.00 | 1.00 |
| nram | 50.00 | 41.84 | 0.00 | 19.39 | 41.84 | 22.45 | 14.29 | 0.00 | 17.35 | 0.00 | 0.00 | 0.00 |
| pa2ga | 37.37 | 20.20 | 2.02 | 20.20 | 35.35 | 2.02 | 17.53 | 1.02 | 9.18 | 0.00 | 0.00 | 0.00 |
| parp1 | 36.22 | 18.50 | 0.98 | 14.17 | 41.12 | 19.09 | 12.40 | 13.78 | 16.14 | 2.56 | 1.18 | 0.79 |
| pde5a | 27.14 | 13.82 | 2.76 | 10.55 | 21.77 | 8.79 | 1.76 | 11.81 | 14.32 | 1.01 | 1.26 | 0.50 |
| pgh1 | 0.00 | 0.00 | 3.08 | 14.36 | 17.11 | 7.18 | 5.21 | 6.15 | 9.23 | 1.54 | 1.03 | 9.23 |
| pgh2 | 0.92 | 0.00 | 5.06 | 18.62 | 34.91 | 14.94 | 11.08 | 25.98 | 17.24 | 1.84 | 1.38 | 12.18 |
| plk1 | 31.78 | 38.32 | 0.00 | 30.84 | 43.81 | 3.74 | 8.41 | 0.00 | 38.68 | 5.66 | 5.66 | 0.93 |
| pnph | 65.05 | 63.11 | 19.42 | 25.24 | 7.77 | 10.68 | 10.68 | 11.76 | 35.29 | 0.00 | 1.96 | 3.88 |
| ppara | 50.13 | 22.25 | 1.34 | 17.43 | 8.94 | 12.60 | 3.77 | 6.70 | 4.83 | 0.80 | 2.14 | 0.80 |
| ppard | 48.75 | 33.75 | 3.33 | 12.08 | 3.61 | 10.42 | 1.25 | 1.25 | 4.58 | 1.25 | 0.42 | 1.25 |
| pparg | 50.00 | 6.61 | 1.24 | 17.36 | 7.53 | 4.96 | 1.66 | 6.20 | 5.99 | 3.10 | 1.24 | 1.03 |
| prgr | 22.87 | 9.90 | 3.41 | 2.39 | 19.05 | 4.44 | 3.64 | 12.97 | 18.43 | 2.39 | 3.75 | 7.85 |
| ptn1 | 24.62 | 18.46 | 0.77 | 41.54 | 15.45 | 10.77 | 14.84 | 26.15 | 25.38 | 10.00 | 4.62 | 0.00 |
| pur2 | 56.00 | 12.00 | 6.00 | 56.00 | 54.00 | 56.00 | 56.00 | 4.00 | 52.00 | 0.00 | 12.00 | 0.00 |
| pygm | 0.00 | 1.30 | 1.30 | 6.49 | 1.30 | 0.00 | 9.09 | 2.60 | 3.90 | 0.00 | 9.09 | 0.00 |
| pyrd | 0.90 | 0.00 | 0.00 | 31.53 | 41.18 | 11.71 | 29.09 | 20.72 | 22.52 | 11.71 | 0.00 | 3.60 |
| reni | 17.31 | 3.85 | 0.00 | 24.04 | 33.98 | 36.54 | 11.65 | 4.85 | 8.74 | 3.88 | 0.97 | 0.00 |
| rock1 | 24.00 | 25.00 | 4.00 | 12.00 | 25.25 | 4.00 | 23.00 | 7.00 | 19.00 | 1.00 | 1.00 | 2.00 |
| rxra | 54.20 | 24.43 | 2.29 | 30.53 | 39.32 | 27.48 | 7.69 | 29.77 | 35.88 | 0.76 | 0.00 | 35.11 |
| sahh | 12.70 | 0.00 | 0.00 | 42.86 | 43.33 | 55.56 | 30.16 | 22.22 | 0.00 | 0.00 | 0.00 | 0.00 |
| src | 59.16 | 50.38 | 1.91 | 14.31 | 12.21 | 7.82 | 14.76 | 5.53 | 11.83 | 1.53 | 1.72 | 13.17 |
| tgfr1 | 54.89 | 31.58 | 10.53 | 38.35 | 44.36 | 18.05 | 25.56 | 10.53 | 28.57 | 0.75 | 0.00 | 1.50 |
| thb | 58.25 | 0.00 | 0.00 | 31.07 | 35.79 | 16.50 | 15.53 | 26.21 | 34.95 | 1.94 | 0.00 | 1.94 |
| thrb | 54.66 | 34.27 | 2.17 | 32.75 | 39.47 | 27.55 | 33.11 | 1.52 | 35.14 | 7.38 | 5.86 | 0.00 |
| try1 | 54.79 | 42.09 | 4.23 | 42.76 | 44.19 | 40.98 | 50.56 | 2.67 | 22.94 | 9.35 | 3.34 | 0.22 |
| tryb1 | 12.84 | 15.54 | 1.35 | 33.78 | 28.57 | 19.59 | 27.89 | 8.11 | 2.03 | 7.43 | 0.68 | 0.00 |
| tysy | 48.62 | 0.00 | 3.67 | 45.87 | 38.53 | 23.85 | 18.35 | 22.94 | 30.28 | 3.67 | 2.75 | 0.00 |
| urok | 53.70 | 48.77 | 2.47 | 43.21 | 53.70 | 12.96 | 47.53 | 8.02 | 12.35 | 2.47 | 0.00 | 3.09 |
| vgfr2 | 37.90 | 36.19 | 4.65 | 20.54 | 17.00 | 4.65 | 31.68 | 17.36 | 22.74 | 5.87 | 0.00 | 3.67 |
| wee1 | 55.88 | 16.67 | 0.00 | 61.76 | 59.80 | 52.94 | 40.20 | 54.46 | 59.41 | 1.98 | 0.00 | 0.98 |
| xiap | 32.00 | 16.00 | 0.00 | 26.00 | 50.51 | 37.00 | 27.00 | 10.10 | 35.35 | 0.00 | 4.04 | 0.00 |

Table S2: DUD-E per-target $EF_1$ scores. Abbreviations as in Table S1.

| | D-G | D-R | DDTA | Gold | Glide | Surflex | Flexx | Vina | GNINA | RF | NN | Bsl. |
|---|---|---|---|---|---|---|---|---|---|---|---|---|
| aa2ar | 0.53 | 0.00 | 0.05 | 0.29 | 0.13 | 0.32 | 0.11 | 0.06 | 0.14 | 0.04 | 0.01 | 0.04 |
| abl1 | 0.73 | 0.63 | 0.12 | 0.46 | 0.33 | 0.30 | 0.48 | 0.25 | 0.34 | 0.19 | 0.03 | 0.14 |
| ace | 0.48 | 0.52 | 0.00 | 0.27 | 0.09 | 0.07 | 0.03 | 0.04 | 0.08 | 0.10 | 0.10 | 0.00 |
| aces | 0.38 | 0.09 | 0.14 | 0.36 | 0.16 | 0.34 | 0.03 | 0.25 | 0.27 | 0.09 | 0.03 | 0.07 |
| ada | 0.85 | 0.67 | 0.03 | 0.32 | 0.15 | 0.18 | 0.20 | 0.02 | 0.23 | 0.10 | 0.00 | 0.00 |
| ada17 | 0.95 | 0.89 | 0.01 | 0.34 | 0.43 | 0.57 | 0.04 | 0.42 | 0.48 | 0.07 | 0.07 | 0.06 |
| adrb1 | 0.38 | 0.01 | 0.04 | 0.45 | 0.32 | 0.26 | 0.18 | 0.08 | 0.08 | 0.03 | 0.03 | 0.01 |
| adrb2 | 0.45 | 0.03 | 0.04 | 0.42 | 0.47 | 0.33 | 0.34 | 0.09 | 0.13 | 0.02 | 0.06 | 0.02 |
| akt1 | 0.36 | 0.58 | 0.10 | 0.38 | 0.21 | 0.08 | 0.10 | 0.13 | 0.11 | 0.04 | 0.01 | 0.03 |
| akt2 | 0.50 | 0.28 | 0.10 | 0.63 | 0.42 | 0.24 | 0.24 | 0.39 | 0.22 | 0.02 | 0.05 | 0.07 |
| aldr | 0.11 | 0.04 | 0.01 | 0.40 | 0.43 | 0.15 | 0.22 | 0.16 | 0.17 | 0.03 | 0.01 | 0.04 |
| ampc | 0.26 | 0.23 | 0.00 | 0.04 | 0.09 | 0.02 | 0.04 | 0.02 | 0.02 | 0.01 | 0.01 | 0.00 |
| andr | 0.60 | 0.38 | 0.02 | 0.04 | 0.42 | 0.08 | 0.01 | 0.31 | 0.17 | 0.03 | 0.04 | 0.05 |
| aofb | 0.05 | 0.02 | 0.02 | 0.27 | 0.14 | 0.10 | 0.11 | 0.16 | 0.04 | 0.02 | 0.02 | 0.07 |
| bace1 | 0.77 | 0.33 | 0.02 | 0.41 | 0.26 | 0.29 | 0.21 | 0.09 | 0.32 | 0.30 | 0.03 | 0.01 |
| braf | 0.71 | 0.67 | 0.09 | 0.39 | 0.50 | 0.10 | 0.41 | 0.27 | 0.34 | 0.20 | 0.03 | 0.05 |
| cah2 | 0.92 | 0.89 | 0.05 | 0.29 | 0.08 | 0.06 | 0.13 | 0.01 | 0.44 | 0.30 | 0.06 | 0.24 |
| casp3 | 0.63 | 0.00 | 0.01 | 0.49 | 0.46 | 0.23 | 0.38 | 0.05 | 0.30 | 0.04 | 0.07 | 0.01 |
| cdk2 | 0.74 | 0.52 | 0.07 | 0.29 | 0.50 | 0.07 | 0.22 | 0.17 | 0.37 | 0.02 | 0.03 | 0.17 |
| comt | 0.86 | 0.08 | 0.00 | 0.82 | 0.95 | 0.05 | 0.15 | 0.06 | 0.02 | 0.00 | 0.06 | 0.00 |
| cp2c9 | 0.17 | 0.09 | 0.03 | 0.12 | 0.04 | 0.05 | 0.17 | 0.05 | 0.08 | 0.09 | 0.01 | 0.01 |
| cp3a4 | 0.05 | 0.06 | 0.05 | 0.22 | 0.17 | 0.14 | 0.07 | 0.04 | 0.03 | 0.07 | 0.01 | 0.06 |
| csf1r | 0.67 | 0.44 | 0.11 | 0.34 | 0.31 | 0.05 | 0.38 | 0.05 | 0.31 | NaN | NaN | 0.07 |
| cxcr4 | 0.43 | 0.14 | 0.03 | 0.11 | 0.01 | 0.26 | 0.01 | 0.00 | 0.33 | 0.00 | 0.10 | 0.01 |
| def | 0.97 | 0.82 | 0.00 | 0.48 | 0.14 | 0.59 | 0.01 | 0.20 | 0.58 | 0.08 | 0.04 | 0.00 |
| dhi1 | 0.00 | 0.05 | 0.02 | 0.16 | 0.13 | 0.09 | 0.01 | 0.09 | 0.20 | 0.12 | 0.04 | 0.03 |
| dpp4 | 0.44 | 0.07 | 0.08 | 0.30 | 0.22 | 0.10 | 0.22 | 0.02 | 0.23 | 0.02 | 0.06 | 0.04 |
| drd3 | 0.12 | 0.02 | 0.02 | 0.18 | 0.04 | 0.13 | 0.05 | 0.10 | 0.14 | 0.08 | 0.03 | 0.08 |
| dyr | 0.80 | 0.69 | 0.07 | 0.66 | 0.47 | 0.40 | 0.17 | 0.12 | 0.44 | 0.03 | 0.04 | 0.04 |
| egfr | 0.87 | 0.70 | 0.05 | 0.40 | 0.41 | 0.22 | 0.31 | 0.10 | 0.35 | 0.06 | 0.06 | 0.14 |
| esr1 | 0.88 | 0.20 | 0.04 | 0.42 | 0.81 | 0.47 | 0.51 | 0.35 | 0.43 | 0.00 | 0.02 | 0.03 |
| esr2 | 0.80 | 0.21 | 0.04 | 0.31 | 0.75 | 0.39 | 0.51 | 0.28 | 0.46 | 0.00 | 0.01 | 0.03 |
| fa10 | 0.70 | 0.62 | 0.11 | 0.72 | 0.31 | 0.24 | 0.72 | 0.38 | 0.61 | 0.31 | 0.03 | 0.06 |
| fa7 | 0.92 | 0.72 | 0.07 | 0.77 | 0.71 | 0.73 | 0.90 | 0.28 | 0.52 | 0.35 | 0.02 | 0.00 |
| fabp4 | 0.77 | 0.33 | 0.00 | 0.50 | 0.48 | 0.10 | 0.33 | 0.43 | 0.01 | 0.05 | 0.00 | 0.00 |
| fak1 | 0.98 | 0.93 | 0.03 | 0.52 | 0.33 | 0.09 | 0.31 | 0.33 | 0.43 | 0.05 | 0.00 | 0.14 |
| fgfr1 | 0.38 | 0.36 | 0.44 | 0.83 | 0.04 | 0.28 | 0.30 | 1.00 | 0.66 | 1.00 | 1.00 | 0.30 |
| fkb1a | 0.55 | 0.26 | 0.00 | 0.39 | 0.71 | 0.19 | 0.01 | 0.13 | 0.09 | 0.03 | 0.01 | 0.03 |
| fnta | 0.67 | 0.05 | 0.02 | 0.32 | 0.12 | 0.13 | 0.11 | 0.05 | 0.13 | 0.11 | 0.03 | 0.03 |
| fpps | 0.01 | 0.02 | 0.00 | 0.97 | 0.01 | 0.03 | 0.07 | 0.00 | 0.52 | 0.00 | 0.92 | 0.00 |
| gcr | 0.16 | 0.27 | 0.04 | 0.13 | 0.25 | 0.26 | 0.13 | 0.24 | 0.29 | 0.05 | 0.06 | 0.07 |
| glcm | 0.01 | 0.07 | 0.03 | 0.40 | 0.18 | 0.49 | 0.16 | 0.00 | 0.10 | 0.00 | 0.05 | 0.00 |

| | | | | | | | | | | | | |
|---|---|---|---|---|---|---|---|---|---|---|---|---|
| gria2 | 0.23 | 0.23 | 0.03 | 0.23 | 0.48 | 0.14 | 0.13 | 0.17 | 0.30 | 0.07 | 0.14 | 0.05 |
| grik1 | 0.63 | 0.48 | 0.03 | 0.47 | 0.33 | 0.18 | 0.38 | 0.08 | 0.20 | 0.03 | 0.06 | 0.03 |
| hdac2 | 0.87 | 0.78 | 0.06 | 0.32 | 0.27 | 0.26 | 0.29 | 0.24 | 0.09 | 0.08 | 0.05 | 0.09 |
| hdac8 | 0.93 | 0.86 | 0.04 | 0.21 | 0.10 | 0.09 | 0.23 | 0.34 | 0.31 | 0.07 | 0.02 | 0.12 |
| hivint | 0.01 | 0.01 | 0.01 | 0.17 | 0.04 | 0.07 | 0.07 | 0.05 | 0.01 | 0.00 | 0.00 | 0.02 |
| hivpr | 0.73 | 0.41 | 0.03 | 0.28 | 0.16 | 0.11 | 0.04 | 0.09 | 0.45 | 0.36 | 0.01 | 0.11 |
| hivrt | 0.31 | 0.05 | 0.01 | 0.23 | 0.41 | 0.09 | 0.05 | 0.09 | 0.22 | 0.02 | 0.05 | 0.10 |
| hmdh | 0.71 | 0.70 | 0.01 | 0.42 | 0.69 | 0.34 | 0.05 | 0.10 | 0.65 | 0.10 | 0.04 | 0.01 |
| hs90a | 0.69 | 0.27 | 0.05 | 0.26 | 0.03 | 0.02 | 0.03 | 0.00 | 0.17 | 0.00 | 0.04 | 0.03 |
| hxk4 | 0.32 | 0.07 | 0.05 | 0.31 | 0.38 | 0.03 | 0.09 | 0.05 | 0.31 | 0.01 | 0.00 | 0.27 |
| igf1r | 0.78 | 0.69 | 0.06 | 0.46 | 0.50 | 0.19 | 0.47 | 0.25 | 0.48 | 0.20 | 0.04 | 0.05 |
| inha | 0.38 | 0.00 | 0.00 | 0.55 | 0.15 | 0.19 | 0.12 | 0.20 | 0.04 | 0.03 | 0.00 | 0.01 |
| ital | 0.69 | 0.20 | 0.00 | 0.12 | 0.13 | 0.04 | 0.03 | 0.01 | 0.11 | 0.05 | 0.01 | 0.06 |
| jak2 | 0.90 | 0.75 | 0.12 | 0.49 | 0.51 | 0.21 | 0.16 | 0.26 | 0.47 | 0.13 | 0.06 | 0.15 |
| kif11 | 0.46 | 0.27 | 0.01 | 0.54 | 0.59 | 0.11 | 0.09 | 0.44 | 0.57 | 0.29 | 0.01 | 0.07 |
| kit | 0.60 | 0.52 | 0.07 | 0.25 | 0.11 | 0.06 | 0.16 | 0.10 | 0.22 | 0.12 | 0.03 | 0.09 |
| kith | 0.00 | 0.01 | 0.00 | 0.30 | 0.63 | 0.70 | 0.08 | 0.51 | 0.61 | 0.06 | 0.04 | 0.00 |
| kpcb | 0.77 | 0.50 | 0.01 | 0.50 | 0.69 | 0.44 | 0.44 | 0.46 | 0.36 | 0.03 | 0.04 | 0.13 |
| lck | 0.74 | 0.67 | 0.07 | 0.26 | 0.39 | 0.22 | 0.31 | 0.16 | 0.26 | 0.07 | 0.02 | 0.06 |
| lkha4 | 0.85 | 0.57 | 0.00 | 0.28 | 0.57 | 0.34 | 0.15 | 0.29 | 0.30 | 0.02 | 0.00 | 0.04 |
| mapk2 | 0.70 | 0.51 | 0.26 | 0.43 | 0.49 | 0.05 | 0.20 | 0.27 | 0.31 | 0.04 | 0.03 | 0.12 |
| mcr | 0.51 | 0.38 | 0.02 | 0.05 | 0.39 | 0.06 | 0.01 | 0.14 | 0.23 | 0.11 | 0.20 | 0.11 |
| met | 0.84 | 0.80 | 0.03 | 0.61 | 0.44 | 0.15 | 0.35 | 0.22 | 0.56 | 0.32 | 0.03 | 0.05 |
| mk01 | 0.87 | 0.74 | 0.05 | 0.44 | 0.28 | 0.04 | 0.38 | 0.11 | 0.19 | 0.11 | 0.06 | 0.02 |
| mk10 | 0.68 | 0.46 | 0.10 | 0.41 | 0.26 | 0.07 | 0.25 | 0.14 | 0.13 | 0.07 | 0.03 | 0.09 |
| mk14 | 0.87 | 0.45 | 0.06 | 0.26 | 0.36 | 0.06 | 0.26 | 0.13 | 0.32 | 0.13 | 0.02 | 0.12 |
| mmp13 | 0.95 | 0.88 | 0.03 | 0.40 | 0.30 | 0.24 | 0.08 | 0.08 | 0.17 | 0.10 | 0.04 | 0.17 |
| mp2k1 | 0.41 | 0.44 | 0.01 | 0.37 | 0.22 | 0.02 | 0.16 | 0.01 | 0.13 | 0.02 | 0.07 | 0.14 |
| nos1 | 0.05 | 0.02 | 0.04 | 0.30 | 0.20 | 0.33 | 0.14 | 0.03 | 0.13 | 0.01 | 0.04 | 0.02 |
| nram | 0.80 | 0.67 | 0.00 | 0.39 | 0.68 | 0.40 | 0.21 | 0.00 | 0.30 | 0.01 | 0.03 | 0.00 |
| pa2ga | 0.66 | 0.43 | 0.04 | 0.34 | 0.59 | 0.06 | 0.32 | 0.02 | 0.16 | 0.03 | 0.02 | 0.01 |
| parp1 | 0.58 | 0.33 | 0.03 | 0.24 | 0.67 | 0.32 | 0.22 | 0.26 | 0.31 | 0.06 | 0.02 | 0.03 |
| pde5a | 0.42 | 0.22 | 0.06 | 0.18 | 0.34 | 0.15 | 0.04 | 0.19 | 0.24 | 0.02 | 0.02 | 0.03 |
| pgh1 | 0.00 | 0.00 | 0.07 | 0.25 | 0.30 | 0.16 | 0.09 | 0.11 | 0.17 | 0.04 | 0.01 | 0.19 |
| pgh2 | 0.02 | 0.01 | 0.11 | 0.33 | 0.60 | 0.28 | 0.22 | 0.45 | 0.29 | 0.04 | 0.02 | 0.24 |
| plk1 | 0.51 | 0.60 | 0.00 | 0.49 | 0.62 | 0.07 | 0.19 | 0.03 | 0.61 | 0.11 | 0.09 | 0.03 |
| pnph | 0.97 | 0.88 | 0.30 | 0.47 | 0.16 | 0.18 | 0.17 | 0.20 | 0.57 | 0.01 | 0.04 | 0.06 |
| ppara | 0.84 | 0.41 | 0.04 | 0.34 | 0.21 | 0.26 | 0.10 | 0.14 | 0.11 | 0.02 | 0.04 | 0.02 |
| ppard | 0.86 | 0.62 | 0.07 | 0.24 | 0.11 | 0.24 | 0.04 | 0.05 | 0.08 | 0.04 | 0.01 | 0.04 |
| pparg | 0.78 | 0.18 | 0.03 | 0.34 | 0.17 | 0.11 | 0.04 | 0.13 | 0.12 | 0.08 | 0.03 | 0.03 |
| prgr | 0.40 | 0.19 | 0.07 | 0.07 | 0.22 | 0.08 | 0.09 | 0.22 | 0.32 | 0.05 | 0.09 | 0.16 |
| ptn1 | 0.42 | 0.28 | 0.03 | 0.66 | 0.34 | 0.23 | 0.28 | 0.45 | 0.45 | 0.20 | 0.08 | 0.04 |
| pur2 | 1.00 | 0.29 | 0.16 | 0.95 | 0.97 | 0.98 | 0.98 | 0.15 | 0.92 | 0.00 | 0.29 | 0.01 |
| pygm | 0.00 | 0.02 | 0.03 | 0.19 | 0.02 | 0.01 | 0.13 | 0.04 | 0.08 | 0.00 | 0.20 | 0.00 |
| pyrd | 0.02 | 0.02 | 0.00 | 0.52 | 0.62 | 0.25 | 0.49 | 0.34 | 0.34 | 0.18 | 0.00 | 0.07 |
| reni | 0.26 | 0.07 | 0.01 | 0.44 | 0.48 | 0.56 | 0.21 | 0.11 | 0.15 | 0.06 | 0.02 | 0.00 |
| rock1 | 0.35 | 0.40 | 0.06 | 0.24 | 0.41 | 0.06 | 0.39 | 0.10 | 0.29 | 0.02 | 0.02 | 0.04 |
| rxra | 0.83 | 0.44 | 0.05 | 0.52 | 0.84 | 0.53 | 0.24 | 0.51 | 0.58 | 0.02 | 0.01 | 0.59 |
| sahh | 0.23 | 0.01 | 0.00 | 0.69 | 0.96 | 0.87 | 0.58 | 0.41 | 0.04 | 0.00 | 0.08 | 0.00 |
| src | 0.82 | 0.76 | 0.06 | 0.24 | 0.23 | 0.14 | 0.24 | 0.09 | 0.27 | 0.04 | 0.04 | 0.21 |
| tgfr1 | 0.82 | 0.50 | 0.14 | 0.60 | 0.67 | 0.29 | 0.43 | 0.23 | 0.47 | 0.02 | 0.00 | 0.03 |
| thb | 0.76 | 0.00 | 0.02 | 0.46 | 0.61 | 0.31 | 0.28 | 0.43 | 0.54 | 0.03 | 0.02 | 0.04 |
| thrb | 0.88 | 0.59 | 0.06 | 0.53 | 0.65 | 0.48 | 0.55 | 0.07 | 0.57 | 0.13 | 0.12 | 0.01 |
| try1 | 0.89 | 0.70 | 0.09 | 0.67 | 0.69 | 0.68 | 0.78 | 0.07 | 0.43 | 0.18 | 0.08 | 0.01 |
| tryb1 | 0.28 | 0.29 | 0.03 | 0.60 | 0.47 | 0.38 | 0.45 | 0.16 | 0.07 | 0.16 | 0.03 | 0.01 |
| tysy | 0.73 | 0.00 | 0.07 | 0.70 | 0.64 | 0.42 | 0.32 | 0.36 | 0.49 | 0.08 | 0.05 | 0.03 |
| urok | 0.80 | 0.73 | 0.06 | 0.66 | 0.79 | 0.23 | 0.70 | 0.15 | 0.25 | 0.05 | 0.02 | 0.08 |
| vgfr2 | 0.59 | 0.60 | 0.08 | 0.35 | 0.30 | 0.08 | 0.50 | 0.28 | 0.38 | 0.12 | 0.01 | 0.08 |
| wee1 | 0.93 | 0.25 | 0.00 | 0.89 | 0.99 | 0.78 | 0.65 | 0.81 | 0.84 | 0.08 | 0.01 | 0.02 |
| xiap | 0.56 | 0.29 | 0.00 | 0.50 | 0.77 | 0.64 | 0.58 | 0.16 | 0.59 | 0.01 | 0.09 | 0.00 |

Table S3: DUD-E per-target $BEDROC_{80.5}$ scores. Abbreviations as in Table S1.

| | D-G | D-R | GNINA | DDTA | Bsl. |
|---|---|---|---|---|---|
| ADRB2 | 0.60 | 0.51 | 0.52 | 0.37 | 0.45 |
| ALDH1 | 0.54 | 0.52 | 0.60 | 0.58 | 0.55 |
| ESR1_ago | 0.66 | 0.62 | 0.71 | 0.66 | 0.61 |
| ESR1_ant | 0.52 | 0.57 | 0.72 | 0.63 | 0.57 |
| FEN1 | 0.58 | 0.51 | 0.55 | 0.49 | 0.54 |
| GBA | 0.46 | 0.40 | 0.71 | 0.55 | 0.51 |
| IDH1 | 0.46 | 0.60 | 0.69 | 0.70 | 0.54 |
| KAT2A | 0.39 | 0.47 | 0.41 | 0.48 | 0.49 |
| MAPK1 | 0.61 | 0.64 | 0.62 | 0.58 | 0.51 |
| MTORC1 | 0.57 | 0.61 | 0.49 | 0.47 | 0.50 |
| OPRK1 | 0.82 | 0.70 | 0.80 | 0.69 | 0.76 |
| PKM2 | 0.49 | 0.53 | 0.61 | 0.57 | 0.51 |
| PPARG | 0.76 | 0.74 | 0.76 | 0.68 | 0.59 |
| TP53 | 0.52 | 0.52 | 0.56 | 0.57 | 0.58 |
| VDR | 0.57 | 0.52 | 0.39 | 0.42 | 0.44 |

Table S4: Lit-PCBA per-target AUROC scores. Abbreviations as in Table S1.

| | D-G | D-R | GNINA | DDTA | Bsl. |
|---|---|---|---|---|---|
| ADRB2 | 25.00 | 0.00 | 0.00 | 0.00 | 0.00 |
| ALDH1 | 1.38 | 1.20 | 1.88 | 1.42 | 0.93 |
| ESR1_ago | 8.33 | 8.33 | 15.38 | 0.00 | 0.00 |
| ESR1_ant | 0.99 | 0.00 | 5.88 | 0.00 | 0.00 |
| FEN1 | 3.26 | 3.53 | 1.90 | 0.27 | 3.26 |
| GBA | 3.03 | 1.21 | 9.04 | 4.22 | 1.82 |
| IDH1 | 0.00 | 5.26 | 12.82 | 5.13 | 2.63 |
| KAT2A | 0.52 | 1.55 | 1.55 | 1.03 | 1.04 |
| MAPK1 | 3.58 | 2.28 | 1.62 | 0.65 | 0.65 |
| MTORC1 | 1.04 | 1.04 | 1.03 | 1.03 | 1.04 |
| OPRK1 | 0.00 | 4.35 | 8.33 | 4.17 | 4.35 |
| PKM2 | 0.55 | 0.55 | 0.55 | 0.55 | 1.47 |
| PPARG | 11.54 | 0.00 | 7.41 | 0.00 | 0.00 |
| TP53 | 1.28 | 0.00 | 1.27 | 2.53 | 3.85 |
| VDR | 4.42 | 2.83 | 0.79 | 1.02 | 0.79 |

Table S5: Lit-PCBA per-target $EF_1$ scores. Abbreviations as in Table S1.

| | D-G | D-R | GNINA | DDTA | Bsl. |
|---|---|---|---|---|---|
| ADRB2 | 0.19 | 0.03 | 0.01 | 0.00 | 0.02 |
| ALDH1 | 0.07 | 0.06 | 0.09 | 0.08 | 0.05 |
| ESR1_ago | 0.10 | 0.07 | 0.12 | 0.02 | 0.01 |
| ESR1_ant | 0.04 | 0.01 | 0.12 | 0.04 | 0.00 |
| FEN1 | 0.04 | 0.04 | 0.02 | 0.01 | 0.04 |
| GBA | 0.03 | 0.02 | 0.10 | 0.03 | 0.02 |
| IDH1 | 0.01 | 0.04 | 0.12 | 0.06 | 0.04 |
| KAT2A | 0.01 | 0.02 | 0.02 | 0.02 | 0.01 |
| MAPK1 | 0.05 | 0.03 | 0.02 | 0.02 | 0.01 |
| MTORC1 | 0.01 | 0.01 | 0.01 | 0.01 | 0.02 |
| OPRK1 | 0.01 | 0.03 | 0.07 | 0.03 | 0.03 |
| PKM2 | 0.01 | 0.01 | 0.01 | 0.01 | 0.02 |
| PPARG | 0.15 | 0.00 | 0.06 | 0.00 | 0.00 |
| TP53 | 0.04 | 0.02 | 0.03 | 0.04 | 0.08 |
| VDR | 0.05 | 0.03 | 0.01 | 0.01 | 0.01 |

Table S6: Lit-PCBA per-target $BEDROC_{80.5}$ scores. Abbreviations as in Table S1.
